# A clade 2.3.4.4b H5N1 virus vaccine that elicits cross-protective antibodies against conserved domains of H5 and N1 glycoproteins

**DOI:** 10.1101/2025.08.14.670375

**Authors:** Eduard Puente-Massaguer, Thales Galdino Andrade, Michael J. Scherm, Kirill Vasilev, Hassanein Abozeid, Alesandra J. Rodriguez, Josh Yueh, Disha Bhavsar, John D. Campbell, Dong Yu, Richard J. Webby, Yoshihiro Kawaoka, Gabriele Neumann, Julianna Han, Andrew B. Ward, Florian Krammer

## Abstract

The continuous evolution and widespread dissemination of highly pathogenic avian influenza (HPAI) H5N1 viruses, particularly clade 2.3.4.4b, pose critical challenges to global pandemic preparedness. In this study, we assessed a low-dose inactivated split virus vaccine derived from clade 2.3.4.4b H5N1, formulated with an Alum/CpG adjuvant, using a preclinical mouse model. This vaccine induced potent humoral and cellular immune responses, generating high titers of cross-reactive antibodies targeting both hemagglutinin (HA) and neuraminidase (NA) glycoproteins across homologous and heterologous H5 clades. The Alum/CpG adjuvant enabled significant antigen dose-sparing while promoting a balanced Th1/Th2 immune profile. Functional analyses demonstrated strong virus neutralization, neuraminidase inhibition, and potent antibody-dependent cellular cytotoxicity activity. Additionally, the vaccine elicited robust antigen-specific CD4^+^ and CD8^+^ T cell responses and effectively controlled viral replication in the lungs, accompanied by reduced lung inflammation. Importantly, vaccinated mice were fully protected against lethal challenges with both the homologous clade 2.3.4.4b and heterologous clade 1 H5N1 viruses, despite low hemagglutination inhibition titers. Electron microscopy polyclonal epitope mapping revealed serum antibodies targeting multiple epitopes on homologous HA and NA, with some cross-reacting to conserved epitopes on heterologous proteins, underscoring broad immune recognition. Collectively, these results highlight the potential of this vaccine candidate to provide broad, multifunctional, and durable immunity against both current and emerging H5N1 threats, supporting its further development for pandemic preparedness.

## 1. Introduction

The ongoing global spread and rapid evolution of highly pathogenic avian influenza (HPAI) H5N1 viruses, particularly those of clade 2.3.4.4b (Figure 1A), are a major threat to both animal and human health ^1^. Since their resurgence, these viruses have demonstrated remarkable geographic expansion, affecting a broad range of avian species and increasingly spilling over into mammals, including humans ^2^. The persistent circulation and genetic diversification of clade 2.3.4.4b H5N1 viruses has led to frequent outbreaks and significant economic losses in the poultry industry, and the emergence of H5N1 in dairy cattle has increased concerns about the potential for further adaptation to human hosts. These trends highlight the urgent need for improved and updated vaccines capable of providing broad and durable protection against emerging H5N1 variants.

**Figure 1.**
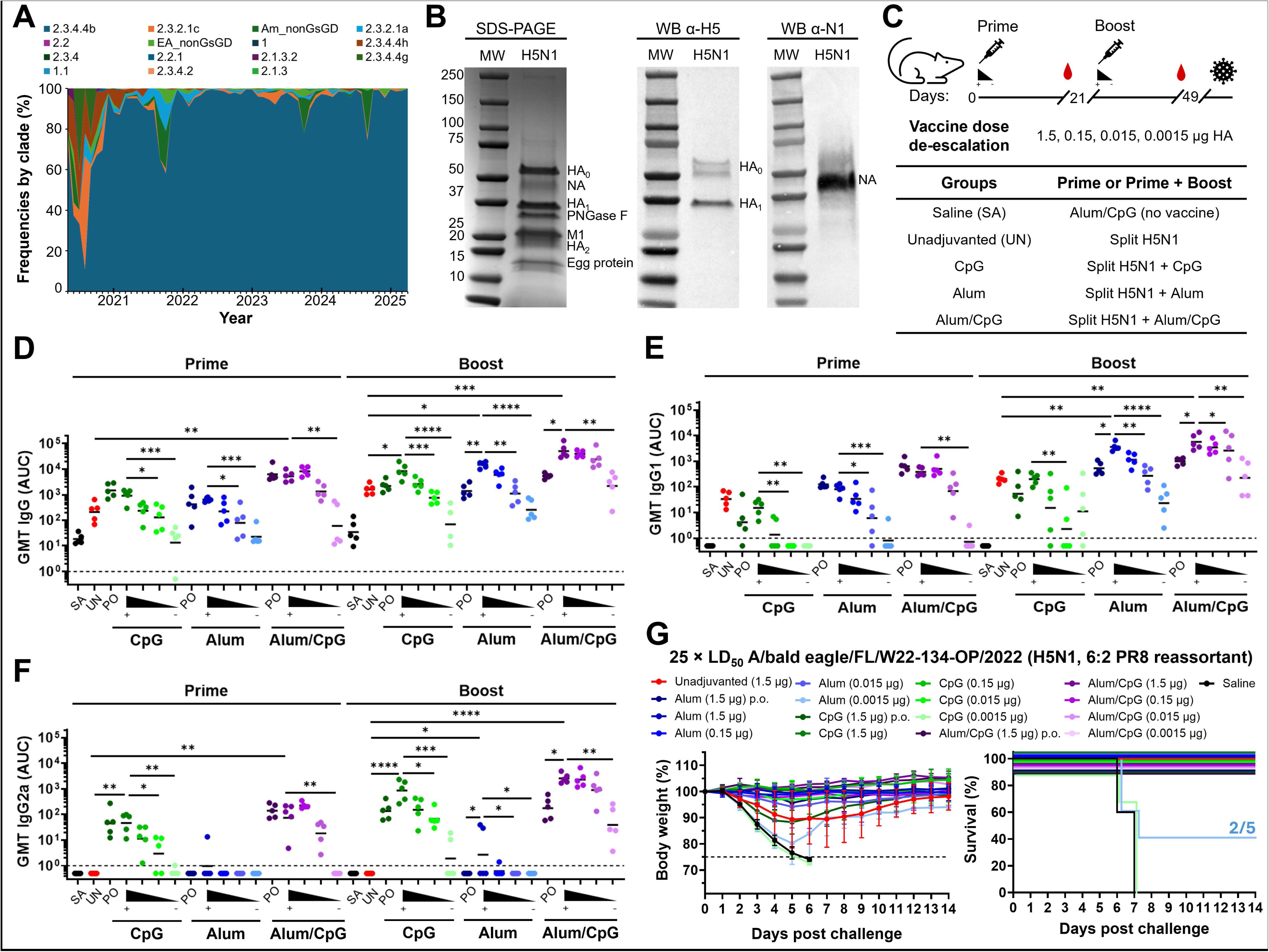
Circulation of H5N1 viruses, vaccine characterization, and vaccine dose-de-escalation in mice. (A) Timeline of global evolutionary frequencies of H5N1 viruses (May-2020 to April-2025). The graph was adapted from Nexstrain.org. (B) Characterization of different vaccine components by SDS-PAGE and confirmation of the presence of H5 and N1 antigens by WB. (C) Experimental design of vaccine dose-de-escalation with and without adjuvants. Different groups of BALB/c mice (*n* = 5) were primed (prime only, p.o.) or prime/boosted (two dose) with different doses of split vaccine (0.0015 – 1.5 μg HA/mouse) adjuvanted with 10 μg/mouse of CpG 1018, 50 μg/mouse of Alum, or both. Vaccine dose-de-escalation was only tested in groups of mice receiving vaccine with an adjuvant in a prime/boost regime. A dose of 1.5 μg HA/mouse was the only condition tested for mice receiving one dose (p.o.) or the unadjuvanted vaccine. All mice were challenged with the A/bald eagle/Florida/W22-134-OP/2022 (H5N1, 6:2 A/PR/8/34) virus (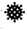) and body weight loss and survival were monitored for 14 days. (D – F) Geometric mean titers (GMT) of total serum IgG, IgG1, and IgG2a responses, respectively, against the closely vaccine matched H5 HA from A/mallard/New York/22-008760-007-original/2022. The black triangle indicates the different vaccine doses assessed from 1.5 to 0.0015 μg HA. Individual and GMT titers are depicted. Statistical comparisons were performed for groups of mice vaccinated with the same vaccine platform (*i.e.* CpG, Alum, or Alum/CpG) including the unadjuvanted vaccine group and excluding the saline group. (G) Percentage body weight loss and Kaplan-Meier survival plots of mice challenged with 25×LD_50_ of the A/bald eagle/Florida/W22-134-OP/2022 (H5N1, 6:2 A/PR/8/34) virus (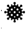). Average body weight values and the standard deviation (SD) are shown. Transverse dotted line denotes the humane endpoint (25% of body weight loss). SA: saline, UN: unadjuvanted, PO: prime only. Only statistically significant *p*-values (<0.05) are shown.

Current pandemic preparedness strategies rely mainly on stockpiled unadjuvanted H5 vaccines or formulated with MF59, AS03 or aluminium-based adjuvants ^3^. While oil-in-water emulsions can enhance antibody responses and offer some cross-clade protection, their efficacy is increasingly undermined by antigenic drift and the static nature of vaccine stockpiles ^4^. As the antigenic landscape of H5N1 continues to shift, there is a growing risk that existing vaccines may provide suboptimal protection against newly emerging clade 2.3.4.4b viruses ^4^. To address these challenges, research is increasingly focused on advancing H5 vaccine design by integrating updated antigens with innovative adjuvant systems that can elicit more potent and broadly protective immune responses. Unlike traditional adjuvants such as MF59 ^5^ and AS03 ^6^ which primarily drive a Th2-skewed response and are generally more reactogenic ^7^, the combination of FDA-approved aluminum hydroxide gels and a toll-like receptor 9 (TLR9) CpG adjuvant can synergistically activate humoral and cellular pathways, fostering a more balanced Th1/Th2 profile and enabling significant antigen dose-sparing ^8,9^.

Importantly, while HA inhibition (HAI) titers remain a well-established correlate of protection for influenza vaccines, accumulating evidence indicates that HAI titers explain only a portion of the overall protective effect. Alternative mechanisms including neutralizing antibodies that target non-HAI epitopes, neuraminidase (NA)-inhibiting antibodies, antibody-dependent cellular cytotoxicity (ADCC), and robust T-cell responses, are increasingly recognized as essential contributors to broad and durable immunity ^10,11^. There is also limited information on the specific epitopes targeted by immune responses elicited by current and experimental pandemic H5 vaccines, particularly on the H5 HA and especially on the N1 NA glycoproteins. Most studies have focused on the magnitude and breadth of antibody responses, with limited information on the epitopes targeted by cross-reactive polyclonal antibody responses ^4,5,12^. This lack of comprehensive epitope mapping restricts our understanding of the mechanisms underlying vaccine-induced protection and impedes the rational design of next-generation vaccines.

In this context, the present study provides a comprehensive evaluation of a clade 2.3.4.4b H5N1 inactivated split virus vaccine formulated with an Alum/CpG adjuvant in a preclinical mouse model. This vaccine was specifically developed to maintain robust immunogenicity against both the H5 HA and N1 NA antigens, thereby aiming for broader protective potential ^13^. Beyond quantifying the magnitude and breadth of antibody responses, this study explores the spectrum of cross-reactive epitopes recognized on both H5 HA and N1 NA glycoproteins across multiple viral clades by negative stain electron microscopy polyclonal epitope mapping (nsEMPEM). This study systematically examines alternative protective mechanisms, including virus neutralization, NAI, ADCC, and the induction of distinct T-cell responses targeting different viral antigens. Through the integration of detailed functional assays with structural analyses, this work not only delineates the complex immunological landscape elicited by the vaccine but also offers critical insights to inform the rational design of next-generation pandemic H5N1 vaccines.

## 2. Results

### 2.1. Low-dose clade 2.3.4.4b H5N1 virus vaccine is highly immunogenic and confers protection against lethal challenge with a homologous virus

An inactivated split virus vaccine (Split) based on the clade 2.3.4.4b genotype B1.1 A/bald eagle/Florida/W22-134-OP/2022 (H5N1, 6:2 A/PR/8/34) was developed using a bioprocess designed to maximize HA and NA antigenicity ^13^. The presence of both H5 and N1 glycoproteins was confirmed by sodium dodecyl-sulphate polyacrylamide gel electrophoresis (SDS-PAGE) and Western blot (WB) (Figure 1B). Different doses of this vaccine (0.0015 – 1.5 μg) were tested in a prime or prime/boost regimen with different adjuvants including CpG 1018^®^ (CpG), aluminum hydroxide gel (Alum), and the combination of both (Alum/CpG, Figure 1C). Prime/boost vaccination resulted in a ∼5 – 10-fold increase in total immunoglobulin G (IgG) titers compared to a single dose of the vaccine (Figure 1D). Combination with Alum/CpG resulted in the highest IgG titers in mouse sera, with similar titers elicited at different vaccine doses (0.015 – 1.5 μg). This vaccine dose-sparing effect was not as pronounced with CpG or Alum alone, especially after boost. Interestingly, 0.015 – 0.15 μg of vaccine with CpG or Alum, or 0.0015 μg with Alum/CpG elicited similar antibody titers to 1.5 μg of unadjuvanted vaccine. Mice in the Alum/CpG group displayed a more balanced Th1 (IgG2a)/Th2 (IgG1) polyclonal IgG response, whereas CpG and Alum alone skewed the immune responses to Th1 and Th2 profiles, respectively (Figure 1E – F).

The protective efficacy of these different vaccine strategies against a highly lethal challenge dose of 25× the median mouse lethal dose (LD_50_) with a homologous virus was assessed (Figure 1G). All mice in the saline group and the lowest vaccine dose (0.0015 μg) with CpG quickly succumbed to the viral infection (100% mortality), and mice vaccinated with the lowest dose with Alum (0.0015 μg) were partially protected from mortality (60% mortality). Mice receiving the unadjuvanted vaccine or a single dose (p.o.) of vaccine (1.5 μg) with CpG were completely protected from mortality but experienced remarkable body weight loss. The rest of mice groups vaccinated with either a single dose or in a prime/boost regimen were fully protected and exhibited little to no body weight loss.

### 2.2. Vaccine combination with diverse adjuvants enhances cross-reactive antibodies against heterologous H5 and N1 glycoproteins and induces durable immunity

Considering the excellent vaccine immunogenicity profile resulting in high IgG titers in sera and complete protection at low doses, 0.15 μg of split vaccine was selected as dose for the following studies (Figure 2A). Sera from mice vaccinated in a prime/boost regimen were analyzed for IgG cross-reactivity against different recombinant HA and NA glycoproteins (Figure 2B). Vaccine combination with adjuvants increased the magnitude and cross-reactivity of IgG responses. This was especially apparent after a single vaccine dose with Alum/CpG. Boosting with an additional dose increased IgG responses in all groups of mice, with Alum/CpG attaining the highest titers followed by Alum, CpG, and the unadjuvanted vaccine. High IgG cross-reactivity titers could also be detected against other clade 2 H5 HAs including clade 2.3.4.4c, the more distant clade 2.3.2.1.c, and even against the clade 1 H5 HA from A/Vietnam/1203/2004. Little cross-reactivity was detected against current seasonal H1 HA or to the stalk (mini HA). Notably, this vaccine induced robust anti-N1 IgG responses across a broad range of N1 NA glycoproteins, targeting not only the matched strain but also phylogenetically distant N1 NAs including those from A/Vietnam/1203/2004 and from recent seasonal strains such as A/Michigan/45/2015 and A/Wisconsin/67/2022.

**Figure 2.**
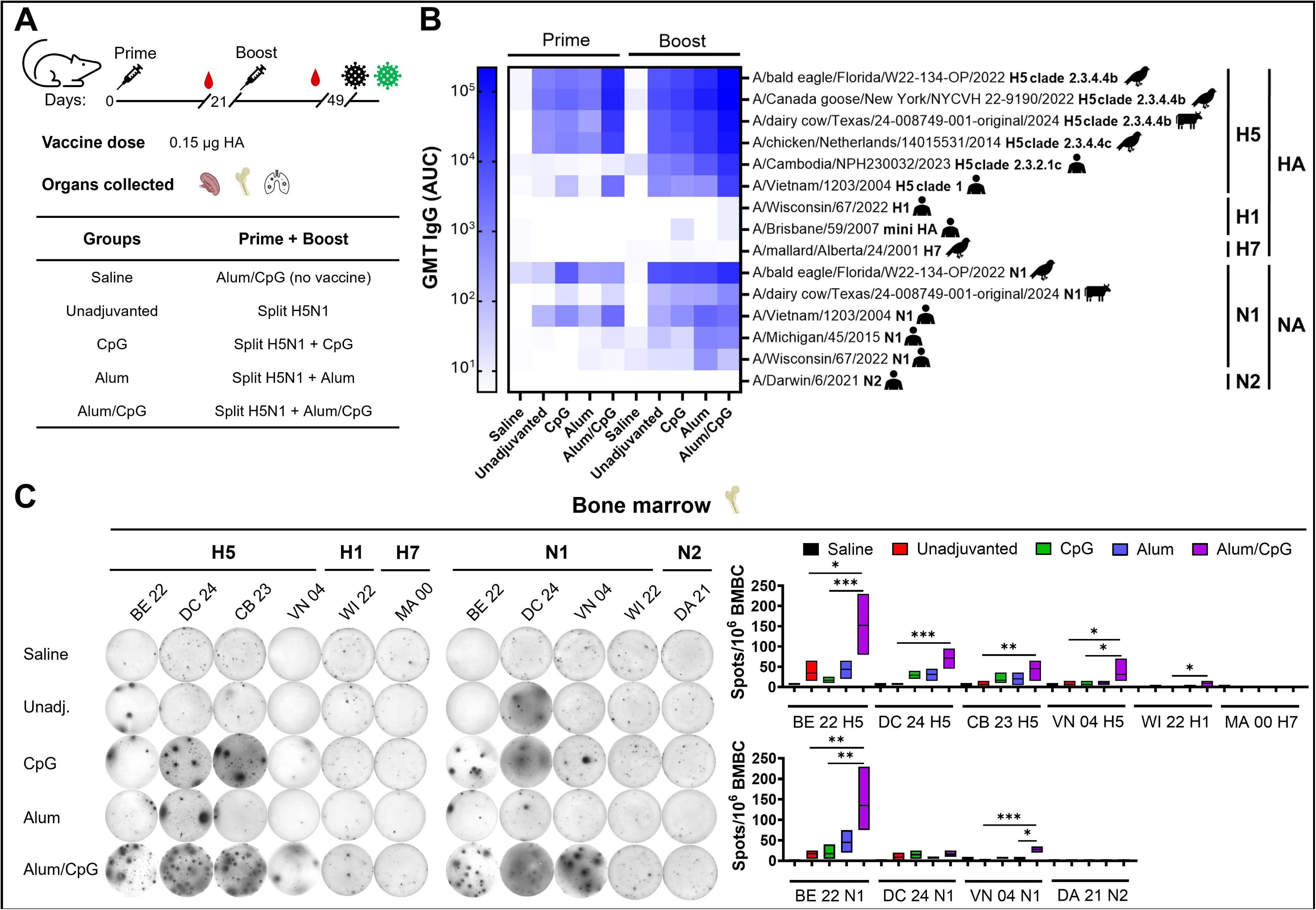
Analysis of antibody cross-reactivity and durability of antigen secreting plasma cell responses. (A) Different groups of BALB/c mice (*n* = 5) were prime/boosted with 0.15 μg HA/mouse of split vaccine adjuvanted with 10 μg/mouse of CpG 1018, 50 μg/mouse of Alum, or both. Spleens, Lungs, and bone marrow were collected, and mice (another set of *n* = 5) was challenged with A/bald eagle/Florida/W22-134-OP/2022 (H5N1, 6:2 A/PR/8/34) virus (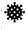) and A/Vietnam/1203/2004 (H5N1, 6:2 A/PR/8/34) virus (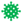). (B) Geometric mean titers (GMT) of total serum IgG responses of mice (*n* = 5) against different HA and NA glycoproteins analyzed by ELISA. (C) Quantification of IgG BMBCs in mice (*n* = 4) at 3 months post-vaccine prime against different HA and NA glycoproteins. One representative mouse per group is shown (left). Box plot (right) with counted spots per 10^6^ BM cells in the ELISpot assay for different HA (top) and NA (bottom) glycoproteins. The average counted spots and SD are shown. Statistical comparisons between groups were conducted for each different glycoprotein excluding the saline group. BE 22: A/bald eagle/FL/W22-134-OP/2022, DC 24: A/dairy cow/Texas/24-008749-001-original/2024, CB 23: A/Cambodia/NPH230032/2023, VN 04: A/Vietnam/1203/2004, WI 22: A/Wisconsin/67/2022, MA 00: A/mallard/Alberta/24/2001, DA 21: A/Darwin/6/2021. Only statistically significant *p*-values (<0.05) are shown.

To analyze the capacity of this vaccine to sustain longer-term cross-reactive H5 and N1 IgG responses, bone marrow antibody secreting B-cells (BMBC) were analyzed 2 – 3 months post vaccination (Figure 2C and Supplementary Figure S1). Mice receiving the unadjuvanted vaccine showed limited induction of cross-reactive BMBCs beyond the homologous H5 HA and N1 NA glycoproteins. A higher number of moderately cross-reactive BMBCs were detected in the Alum and CpG groups, but persistence of cross-reactive BMBCs against the most phylogenetically distant HA and NA antigens was only detected in mice that received the split vaccine in combination with Alum/CpG.

### 2.3. Enhanced antibody and cellular immune functions lead to decreased viral replication and inflammation

Analysis of HAI responses in mice sera against the homologous and a related clade 2.3.4.4b genotype B3.13 (dairy cow) virus isolate indicated that vaccine prime was not sufficient to elicit HAI antibodies. Low HAI titers were detected after boosting in 80% of the mice in the Alum/CpG (Figure 3A, C). However, neutralizing antibodies titers against the homologous virus were detected after prime in all mice receiving an adjuvanted vaccine, and subsequently boosted by ∼3 – 4-fold after the second vaccine dose (Figure 3B). Cross-neutralizing antibody titers against the dairy cow virus isolate were the highest in the Alum/CpG group (Figure 3D). Almost all mice (9/10) in the Alum/CpG group exhibited high NAI antibody titers after prime in comparison to the other groups. After boost, all mice in the Alum and Alum/CpG groups showed the highest NAI antibody titers (Figure 3E). Similarly, antibodies in the sera of mice vaccinated with Alum/CpG displayed the highest level of ADCC *in vitro*, while little to no ADCC activity could be detected in the unadjuvanted vaccine group (Figure 3F).

**Figure 3.**
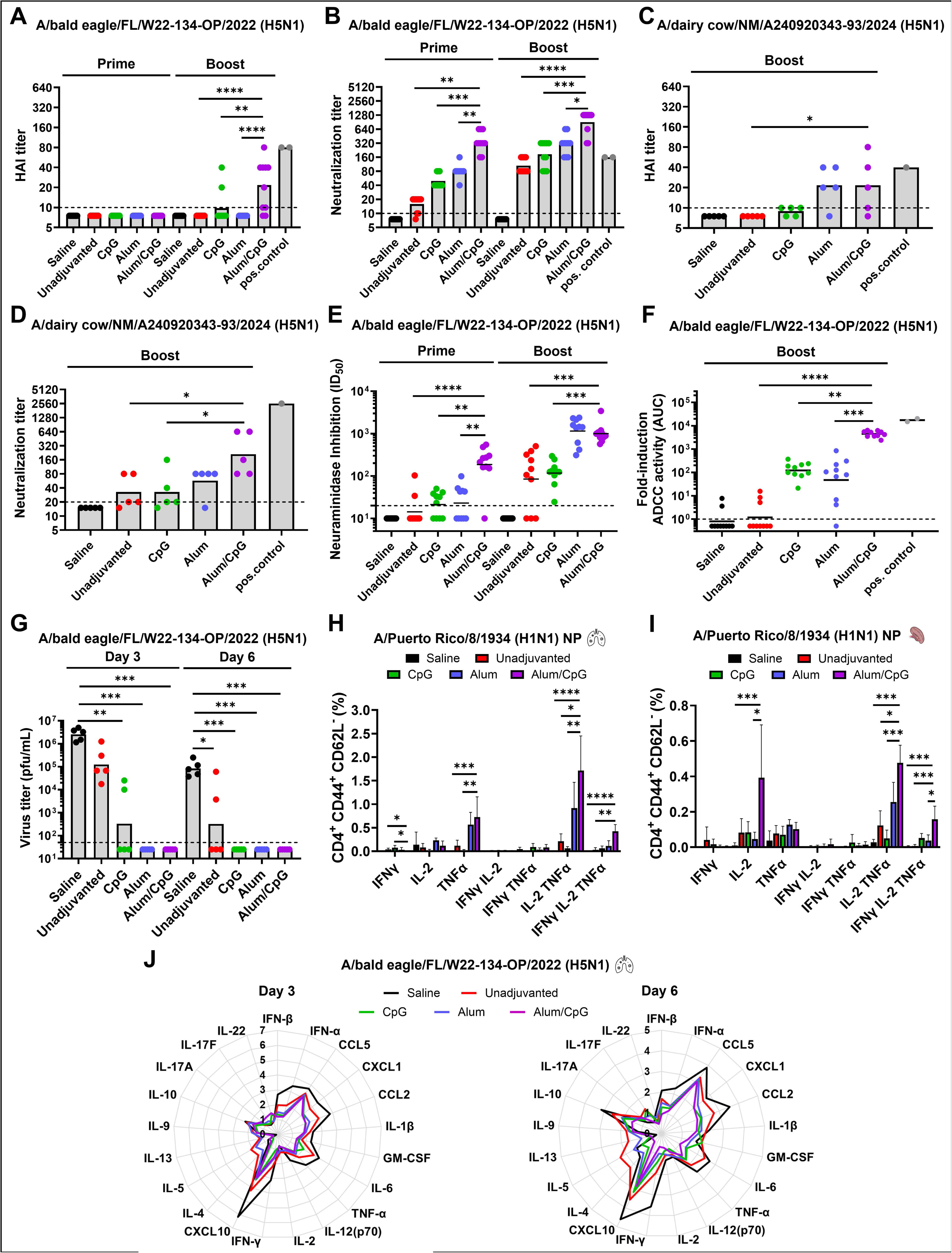
Assessment of vaccine-mediated antibody functions, cellular immunity, and viral replication in lungs. (A – B) HAI and neutralizing activity of mouse (*n* = 10) sera were evaluated against A/bald eagle/Florida/W22-134-OP/2022 (H5N1, 6:2 A/PR/8/34) virus and (C – D) against A/dairy cow/New Mexico/A240920343-93/2024 (H5N1) virus isolate. A *n* = 5 serum samples after vaccine boost was randomly selected to measure HAI and MNT against the A/dairy cow/New Mexico/A240920343-93/2024 (H5N1) virus isolate in BSL-3 conditions. The positive controls used in theses assays include the anti-A/bald eagle/Florida/W22-134-OP/2022 head mAb 1A1 ^36^ (30 μg/mL) in the HAI, mouse anti-A/bald eagle/Florida/W22-134-OP/2022 sera for the MNT assay, and ferret antisera for the HAI and MNT assay against A/dairy cow/New Mexico/A240920343-93/2024. Individual and GMT (grey bars) values for HAI and MNT titers are shown. (E) NA inhibition (NAI) assay of sera (*n* = 10) against A/bald eagle/Florida/W22-134-OP/2022 (H5N1, 6:2 A/PR/8/34) virus. Serum dilutions inhibiting 50% of the NA activity (ID_50_) were plotted. Individual and geometric mean ID_50_ values are shown. (F) Serum of vaccinated mice (*n* = 10) was analyzed for antibody dependent cell mediated cytotoxicity (ADCC) activity against the A/bald eagle/Florida/W22-134-OP/2022 (H5N1, 6:2 A/PR/8/34) virus using a reporter assay. The fold induction of the reporter signal from individual mouse serum over those from blanks were analyzed and plotted as individual and GMT values. (G) Viral load in the lungs of mice vaccinated following a prime/boost regimen and challenged with 0.5×LD_50_ of the A/bald eagle/Florida/W22-134-OP/2022 (H5N1, 6:2 A/PR/8/34) virus. Lungs of BALB/c mice were collected on day 3 (*n* = 5) and 6 (*n* = 5) post challenge. The limit of detection was defined as 50 PFU/mL. A titer of 25 PFU/mL was assigned to negative samples. Individual and GMT (grey bars) values for viral loads are shown (H – I) Percentage of nucleoprotein (NP)-specific CD4^+^ effector memory T-cells in the lungs and spleen of mice (*n* = 5) vaccinated in a prime/boost regimen and challenged with 0.5×LD_50_ of the A/bald eagle/Florida/W22-134-OP/2022 (H5N1, 6:2 A/PR/8/34) virus. Lungs (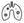) and spleens (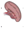) of BALB/c mice were collected on day 5 post challenge. The average T cell percentages and SD are depicted (J). Star plot with mean concentrations of cytokines in mouse lungs on day 3 (*n* = 5) and 6 (*n* = 5) post challenge with 0.5×LD_50_ of the A/bald eagle/Florida/W22-134-OP/2022 (H5N1, 6:2 A/PR/8/34) virus. Statistical comparisons between groups were conducted excluding the saline group except for virus titers and T-cell analysis. Only statistically significant *p*-values (<0.05) are shown. Pos. control: positive control.

Viral replication in the lungs was effectively controlled in all groups receiving vaccines combined with Alum, CpG, or Alum/CpG, with minimal to no detectable virus even at the earliest infection stages (Figure 3G). This control was accompanied by robust CD4^+^ T cell responses targeting NP, H5 HA, and N1 NA, particularly in the Alum/CpG group (Figure 3H–I, Supplementary Figure S2), along with a marked increase in H5 HA-specific CD8^+^ T cells in the lungs of the same group (Supplementary Figure S3). Mice receiving adjuvanted vaccines, especially Alum/CpG, also exhibited reductions in lung inflammation after infection, as evidenced by lower levels of interleukin (IL)-6, interferon (IFN)-α, IFN-γ, tumor necrosis factor (TNF)-α, C-X-C motif chemokine ligand 10 (CXCL10), and other pro-inflammatory mediators compared with saline and unadjuvanted controls, which showed persistently high cytokine levels from early (day 3) to later stages (day 6) of infection (Figure 3J). Notably, the unadjuvanted split vaccine, and to a lesser degree the Alum formulation, induced partial Th2 polarization characterized by IL-4, IL-5, and IL-13 production, whereas the Alum and Alum/CpG groups showed a shift toward a Th17 phenotype with increased IL-22 and IL-17. These immunological differences between adjuvant groups were most evident during the early phase of infection (Figure 3J, Supplementary Figure S4).

### 2.4. Mice vaccinated with a clade 2.3.4.4b H5N1 vaccine in combination with Alum/CpG are fully protected against lethal challenge with a heterologous clade 1 virus

The vaccine’s capacity to elicit diverse antibody responses and confer protection against phylogenetically distant H5N1 viruses was evaluated using the clade 1 A/Vietnam/1203/2004 (H5N1, 6:2 A/PR/8/34) virus (Figure 4A, green). HAI titers were undetectable after two doses of the split vaccine, regardless of adjuvant formulation (Figure 4B). Nevertheless, low levels of neutralizing antibodies were observed post-prime in sera from all mice in the Alum and Alum/CpG groups. All mice developed neutralizing antibodies following two 0.15 μg HA doses of split vaccine, with the highest titers seen in the Alum and Alum/CpG groups (Figure 4C). After priming, 60% of mice in the Alum/CpG group developed NAI antibodies, a response not detected in the other groups. Booster immunization led to a 10-fold increase in NAI titers in the Alum/CpG group, with all Alum-vaccinated mice also seroconverting (Figure 4D). Notably, 50% of the Alum/CpG group also developed NAI antibodies against the more genetically divergent seasonal N1 NA from A/Michigan/45/2015 (Supplementary Figure S5). Sera from mice vaccinated with CpG and Alum/CpG exhibited the strongest ADCC activity *in vitro* against A/Vietnam/1203/2004, with lower ADCC in the Alum group and minimal activity in the unadjuvanted group (Figure 4E).

**Figure 4.**
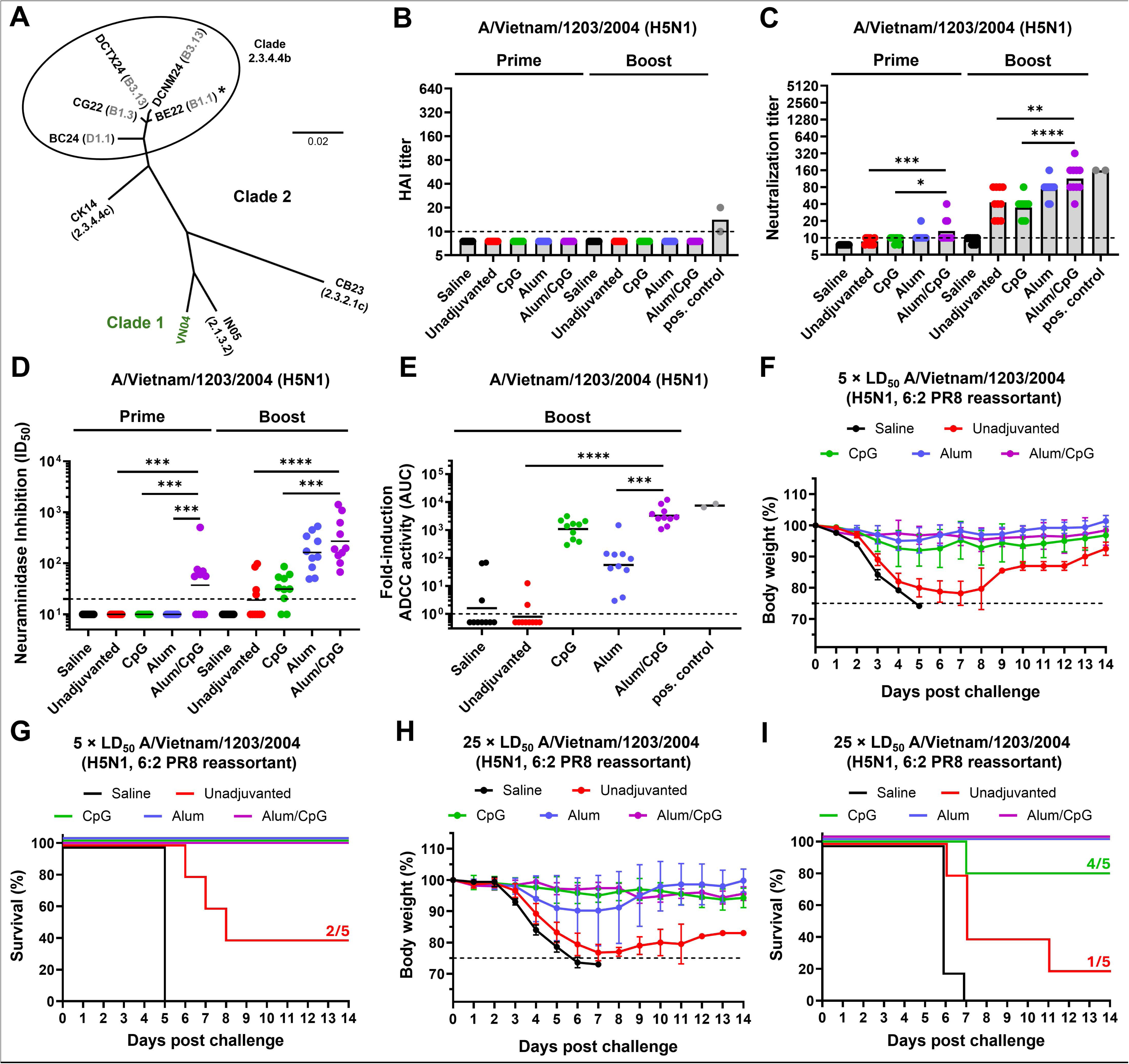
Cross-clade antibody functions and protection against a phylogenetically distant H5N1 virus. (A) Cladogram of HAs from representative H5N1 influenza viruses. The HA from the clade 2.3.4.4b vaccine strain is highlighted with an asterisk. The HA from the clade 1 A/Vietnam/1203/2004 (H5N1, VN04) virus is highlighted in green. The tree was constructed using amino acid sequences aligned in Clustal Omega and visualized with FigTree. BE22: A/bald eagle/Florida/W22-134-OP/2022, DCNM24: A/dairy cow/New Mexico/A240920343-93/2024, DCTX24: A/dairy cow/Texas/24-008749-001-original/2024, CG22: A/Canada goose/New York/NYCVH 22-9190/2022, BC24: A/British Columbia/PHL-2032/2024, CK14: A/chicken/Netherlands/14015531/2014, IN05: A/Indonesia/05/2005, CB23: A/Cambodia/NPH230032/2023. (B – C) HAI and neutralizing activity of mouse (*n* = 10) sera were evaluated against A/Vietnam/1203/2004 (H5N1, 6:2 A/PR/8/34) virus. A mouse anti-A/Vietnam/1203/2004 serum was used as the positive control for the HAI and MNT assays. Individual and GMT (grey bars) values for HAI and MNT titers are shown. (D) NAI assay of sera (*n* = 10) against A/Vietnam/1203/2004 (H5N1, 6:2 A/PR/8/34) virus. Individual and geometric mean ID_50_ values are shown. (E) Serum of vaccinated mice (*n* = 10) was analyzed for ADCC activity against the A/Vietnam/1203/2004 (H5N1, 6:2 A/PR/8/34) virus using a reporter assay. The fold induction of the reporter signal from individual mouse serum over those from blanks were analyzed and plotted as individual and GMT values. (F – I) Percentage body weight loss and Kaplan-Meier survival plots of mice challenged with 5×LD_50_ and 25×LD_50_ of the A/Vietnam/1203/2004 (H5N1, 6:2 A/PR/8/34) virus ( ). Average body weight values and SD are shown. Transverse dotted line denotes the humane endpoint (25% of body weight loss). Statistical comparisons between groups were conducted excluding the saline group. Only statistically significant *p*-values (<0.05) are shown. Pos. control: positive control.

Challenge with a lethal dose (5× LD50) of the clade 1 A/Vietnam/1203/2004 (H5N1, 6:2 A/PR/8/34) virus resulted in complete protection and little to no morbidity in mice vaccinated with the split vaccine in combination with Alum, CpG, and Alum/CpG. Mice receiving saline or the unadjuvanted vaccine either succumbed to the infection or were partially protected (Figure 4F – G). To assess further the protection capacity of this vaccine, a new set of mice was challenged with a 5-fold higher lethal dose (25× LD50) with the same virus. Mice in the Alum/CpG and Alum groups were fully protected from mortality, but 3/5 mice in the Alum group lost substantial body weight while no sign of morbidity was observed in mice vaccinated with Alum/CpG. Eighty percent of the mice in the CpG group were protected from mortality and exhibited low signs of morbidity. Mice vaccinated with saline or the unadjuvanted vaccine succumbed to the infection or were barely protected (1/5), respectively (Figure 4H – I).

### 2.5. Epitope mapping identifies conserved sites on H5 and N1 glycoproteins as targets of cross-reactive antibodies

To dissect epitopes targeted by the polyclonal antibody (pAb) response to H5N1 vaccination, we performed nsEMPEM on serum pooled from five mice per group at 3 – 4 weeks after the second boost. For the vaccine strain-matched A/bald eagle/Florida/W22-134-OP/2022 H5 HA, we observed HA-specific pAbs in the 2D class averages and reconstructed pAb complexes in 3D for all evaluated groups (Figure 5A, Supplementary Figure S6). All the vaccinated groups showed pAbs targeting the HA head, specifically the side of the head, non-receptor binding site (RBS) upper head and the vestigial esterase epitopes (Figure 5B). No response was detected in the saline group. When complexed with NA, the vaccinated groups showed pAbs targeting the top, side, and underside regions on NA (Figure 5C). For A/Vietnam/1203/2004, 2D class averages showed HA stem-specific pAbs in all adjuvanted groups (Figure 5A, Supplementary Figure S6). Polyclonal antibodies targeting the vestigial esterase epitope were also observed in the Alum adjuvant group. No HA response was detected in the saline or unadjuvanted group (Figure 5B). When complexed with NA, the vaccinated groups showed pAbs targeting top and side regions on NA (Figure 5C). For the particle distribution profiles of the nsEMPEM, responses to the central stem and anchor epitopes were grouped as “Stem” and responses to the non-RBS upper head, side head, and the vestigial esterase epitope were grouped as “Head”. For A/bald eagle/Florida/W22-134-OP/2022, we observed a robust response to the HA head, with no unbound antigen being detected in the Alum and Alum/CpG groups (Figure 5D). A similar pattern was observed for the NA, with greater number of fab-antigen complexes in the Alum and Alum/CpG groups, showing 92.2% and 76.8% bound NA, respectively (Figure 5E). For A/Vietnam/1203/2004, fewer pAbs targeting HA and NA were observed compared to the vaccine strain-matched antigens. When comparing particle distributions between HA and NA antigens from each strain, A/bald eagle/Florida/W22-134-OP/2022 showed higher percentages of HA-pAb complexes than NA-pAb. Interestingly, the opposite was observed for A/Vietnam/1203/2004, more pAbs reacted to the NA than to the HA (Figures 5 D – E). Overall, pAbs targeted multiple head epitopes on the homologous HA in high abundance but targeted the stem of heterologous HA in low abundance, while pAbs recognizing homologous and heterologous NAs targeted the top and side regions on NA.

**Figure 5.**
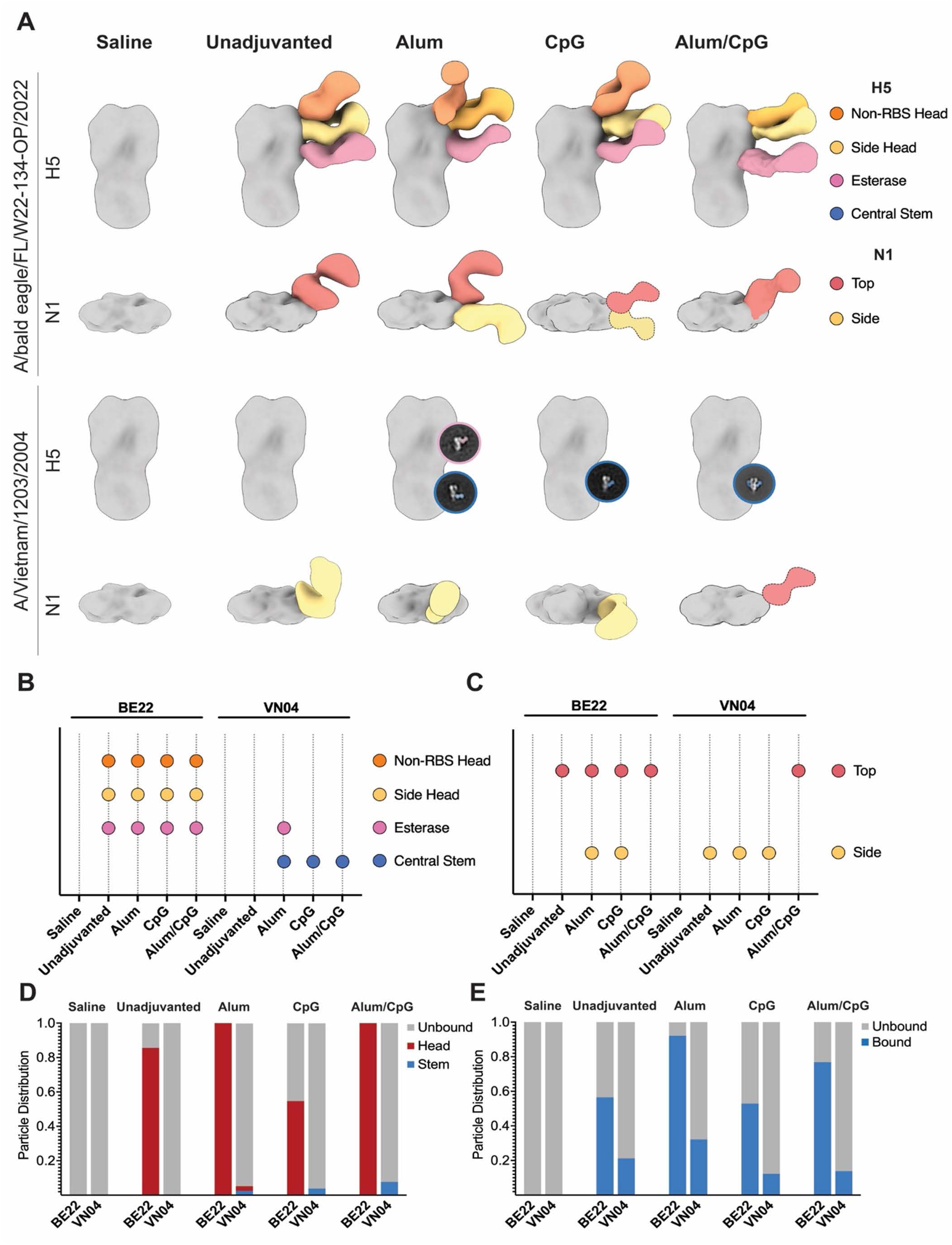
nsEMPEM analysis of polyclonal IgG fab responses in mice 3 – 4 weeks after second boost. (A) Negative stain electron microscopy reconstructions of purified polyclonal IgG fab bound to A/bald eagle/Florida/W22-134-OP/2022 H5 and N1 and A/Vietnam/1203/2004 H5 and N1 glycoproteins. Representative, false-colored 2D classes are presented for epitopes that could not be reconstructed. 3D reconstructions that illustrated epitopes targeted but were poorly resolved are presented with cartoon Fabs. (B – C) Summary of epitopes targeted by pAbs against the H5 HA (B) and N1 NA (C) of A/bald eagle/Florida/W22-134-OP/2022 and A/Vietnam/1203/2004. (D – E) Particle distribution bar charts of free antigen or immune complexed particles targeting HA (D) or NA (E) observed by nsEMPEM. BE22: A/bald eagle/Florida/W22-134-OP/2022, VN04: A/Vietnam/1203/2004.

## 3. Discussion

The present study demonstrates that an inactivated split virus vaccine based on clade 2.3.4.4b H5N1, when formulated with optimized adjuvant combinations, is highly immunogenic and confers robust protection against both homologous and heterologous lethal influenza challenges in mice. The combination of Alum and CpG as adjuvants enabled significant dose-sparing, with as little as 0.0015 μg of vaccine antigen eliciting IgG titers comparable to much higher doses of unadjuvanted vaccine. The Alum/CpG combination not only increased the magnitude of the antibody response but also promoted a balanced Th1/Th2 profile, as evidenced by the IgG2a/IgG1 ratio, in line with the ability to synergistically activate both humoral and cellular arms of the immune system, a feature that has been shown to be critical for broad and durable protection against rapidly evolving influenza viruses ^14^.

A key strength of this vaccine formulation is its ability to induce cross-reactive antibodies against a range of H5 and N1 antigens, including those from phylogenetically distant clades such as 2.3.2.1c (A/Cambodia/NPH230032/2023) ^15^ and clade 1 (A/Vietnam/1203/2004). The Alum/CpG vaccine elicited the highest titers of cross-reactive IgG and, importantly, only this group showed persistent bone marrow-resident antibody-secreting cells against the most divergent HA and NA antigens months after vaccination. The robust anti-N1 response is particularly noteworthy since NA is increasingly recognized as a critical target for cross-protective immunity against influenza A viruses ^16,17^. The ability of the vaccine to elicit anti-N1 antibodies that cross-react with both homologous and heterologous N1 antigens, including those from clade 1 viruses, suggests that this formulation could provide a broader layer of protection, potentially limiting viral replication and transmission even in the face of antigenic drift and shift in the HA and NA glycoproteins.

The Alum/CpG vaccine induced not only high titers of binding antibodies but also functional responses, including neutralizing antibodies detected after a single dose and further boosted after the second dose, with the highest titers in the Alum/CpG group. NAI antibodies were rapidly induced and maintained at high levels, especially in the Alum/CpG and Alum groups. Sera from mice vaccinated with Alum/CpG showed the highest ADCC activity, a mechanism increasingly recognized as important for heterosubtypic protection ^18^. These humoral responses were shown to be durable in the bone marrow and complemented by strong cellular immunity, with increased frequencies of antigen-specific CD4^+^ and CD8^+^ T cells, particularly in the lungs. The Alum/CpG group exhibited the most robust T cell responses against HA, NA, and NP antigens, which likely contributed to the rapid control of viral replication and reduced lung inflammation following challenge. This is in line with recent work showing that broad T cell responses are associated with cross-protection against diverse influenza virus strains ^19^. Remarkably, the Alum/CpG vaccine conferred complete protection against lethal challenge with a heterologous clade 1 H5N1 virus, even in the absence of detectable HAI titers. This robust protection was associated with the induction of cross-neutralizing and cross-NAI antibodies ^20^, as well as strong ADCC activity, highlighting the critical role of non-HAI immune mechanisms in mediating cross-clade immunity ^21^. A key innovation of this study is the structural mapping of pAb serum responses using nsEMPEM, which revealed that vaccination, particularly with Alum/CpG adjuvant, elicited antibodies targeting conserved regions of both H5 HA and N1 NA glycoproteins. Within HA, responses focused on the vestigial esterase site ^22,23^, a critical and relatively conserved region among H5 strains that mediates cross-neutralization despite being generally non-HAI active, as well as other non-HAI epitopes such as the side head and non-RBS head domains. Although epitope mapping of pAb responses to the A/Vietnam/1203/2004 H5 HA was limited likely due to low-affinity or low-resolution fab-antigen interactions, cross-reactive antibodies against the central stem were consistently detected across all adjuvanted vaccine groups, potentially explaining the observed cross-neutralizing non-HAI activity ^24,25^. In the NA, multiple epitopes were identified, including near the catalytic site and several along the lateral face. Notably, epitopes along the NA side region dominated cross-reactive recognition against the A/Vietnam/1203/2004 N1 NA, aligning with monoclonal antibody studies demonstrating that these conserved lateral-face epitopes confer broad NAI and *in vivo* protection ^17,26^. Antibodies targeting N1 NA may also have contributed to the high neutralization titers by limiting viral spread.

The targeting of conserved epitopes within both the HA head and the NA by vaccine-induced antibodies further underscores the potential of this vaccine platform to provide broad and durable immunity against a spectrum of emerging H5Nx and HxN1 viruses. This breadth of protection is especially significant considering the ongoing evolution and geographic expansion of clade 2.3.4.4b H5N1 viruses, which have recently spilled over into mammals and humans, raising concerns about pandemic risk. Consequently, the inclusion of the N1 NA component alongside HA, rather than focusing solely on the H5 HA as in many current vaccine strategies ^27,28^, may offer superior control of pandemic influenza virus transmission and disease by leveraging the complementary protective roles of both HA- and NA-specific antibodies.

Despite these advances, this study has several limitations. All experiments were conducted in mice, necessitating further validation in more predictive models such as ferrets or non-human primates to confirm the durability and breadth of protection. Moreover, while the study highlights several mechanisms of protection, the individual contributions of NAI, ADCC, T cell responses, and antibodies targeting the specific nsEMPEM-identified epitopes remain to be fully disentangled. Future work should also evaluate the impact of pre-existing immunity on vaccine efficacy and assess how rapidly this adjuvant platform can be adapted to counter emerging pandemic influenza threats. In summary, this study demonstrates that a clade 2.3.4.4b H5N1 split vaccine, particularly when adjuvanted with Alum/CpG, is highly immunogenic and amenable to dose-sparing strategies. It elicits broad, durable, and multifunctional immune responses that extend well beyond traditional HAI activity. By providing a comprehensive structural and functional roadmap, this work underscores the importance of targeting conserved epitopes on both HA and NA glycoproteins, offering valuable insights for the rational design of next-generation pandemic influenza vaccines.

## 4. Methods

### 4.1. Cell lines

Baculovirus generation and amplification was performed in Sf9 cells (CRL-1771, ATCC), grown in *Trichoplusia ni* medium-formulation Hink insect cell medium (TNM-FH, Gemini Bioproducts) supplemented with 10% v/v fetal bovine serum (FBS, Gibco), penicillin (100 U/mL) -streptomycin (100 μg/mL) solution (Gibco), and 0.1% v/v Pluronic F-68 (Gibco). High Five cells (BTI-*TN*-5B1-4, B85502, Thermo Fisher Scientific) were grown in Sf900 II SFM (Gibco) and used for recombinant HA and NA production. Both cell lines were grown at 27 °C. HEK293F cells (Thermo Fisher Scientific) cultured at 37°C, 8% CO_2_, shaken at 125 rpm in FreeStyle 293 expression medium (Gibco) and used for recombinant protein production.

Madin-Darby canine kidney (MDCK) cells were grown in modified Eagle’s medium (MEM) containing 10% v/v FBS and penicillin (100 U/mL) -streptomycin (100 μg/mL) solution in a humidified incubator at 37 °C and 5% CO_2_.

### 4.2. Recombinant proteins

Recombinant HA and NA proteins were produced in High Five cells ^29^, and the N1 from A/Vietnam/1203/2004 was expressed and purified from transiently transfected HEK293F cells ^30^. Cell culture supernatants were purified using a Ni^2+^ -nitrilotriacetic acid agarose beads (Qiagen) after incubation at room temperature (RT) for 2 h or overnight (O/N) at 4 °C. Samples were subsequently run over a gravity flow column (Qiagen) and washed with phosphate-buffered saline (PBS) or tris-buffered saline (TBS) and 20 mM imidazole, pH 8.0. Proteins were eluted with 300 – 500 mM imidazole and buffer-exchanged into PBS or TBS three times on a 30-kDa concentrator (Amicon). Affinity-purified N1 from A/Vietnam/1203/2004 was further processed using size exclusion chromatography over a Superdex 200 Increase 10/300 column (Cytiva). Fractions corresponding to tetrameric NA were pooled, concentrated, and buffer-exchanged to TBS using a 30-kDa Amicon concentrator.

### 4.3. Viruses

A/bald eagle/Florida/W22-134-OP/2022 (H5N1, 6:2 A/PR/8/34) ^31^, A/Vietnam/1203/2004 (H5N1, 6:2 A/PR/8/34) and H7N1_A/Michigan/45/2015_ viruses were generated by reverse genetics. The H5 and N1 glycoproteins were derived from the wildtype A/bald eagle/Florida/W22-134-OP/2022 (H5N1) and A/Vietnam/1203/2004 (H5N1) viruses, with removal of the H5 polybasic cleavage site. The H7 and N1 from the H7N1_A/Michigan/45/2015_ virus were derived from A/Shanghai/1/2013 (H7N9) and A/Michigan/45/2015 (H1N1) viruses. The internal gene segments of these three viruses belong to the donor vaccine strain A/Puerto Rico/8/1934 (H1N1, A/PR/8/34). The A/dairy cow/New Mexico/A240920343-93/2024 (H5N1) was isolated in MDCK cells from a milk sample provided by the Texas A&M Veterinary Medical Diagnostic Laboratory ^32^.

The A/bald eagle/Florida/W22-134-OP/2022 (H5N1, 6:2 A/PR/8/34), A/Vietnam/1203/2004 (H5N1, 6:2 A/PR/8/34) and H7N1_A/Michigan/45/2015_ viruses were grown in 10-day old embryonated chicken eggs at 37 °C for 48 h, and cooled at 4 °C O/N. The A/dairy cow/New Mexico/A240920343-93/2024 (H5N1) was grown on MDCK cells. Cell debris was removed by low-speed centrifugation (4000×*g*, 4 °C, 20 min). Viruses were aliquoted, stored at −80 °C, and titrated by the plaque assay method on MDCK cells ^33^.

### 4.4. Production of inactivated split influenza virus vaccines

A/bald eagle/Florida/W22-134-OP/2022 (H5N1, 6:2 A/PR/8/34) vaccine production was conducted as previously described ^13^. Virus inactivation was performed with 0.025% v/v beta-propiolactone (Millipore Sigma) prepared in ice-cold water for injection (Gibco) for 30 min after pH buffering with 0.01 M disodium hydrogen phosphate (Millipore Sigma) and stopped by incubation at 37 °C for 1 h. Then, the inactivated virus sample was centrifuged at 4000 rpm, 4 °C for 30 min. The clarified supernatant was loaded on 5 mL of 30% w/v sucrose cushion prepared in 1X NTE buffer consisting of 1 M NaCl, 100 mM Tris-HCl, 10 mM ethylenediaminetetraacetic acid (EDTA) in water for injection with the pH adjusted to 7.4. The supernatant containing the inactivated virus was concentrated by high-speed centrifugation (25000 rpm, 4 °C for 2 h), and the pelleted virus was resuspended in TBS (pH 7.5). The resuspended virus was split with 1% v/v Triton X-100 (Fisher Bioreagents), and the detergent was removed by incubation with 0.2 g of Bio-Beads SM-2 (BioRad) per mL of inactivated split virus. The supernatant was collected, and the total protein concentration was adjusted to 0.5 mg/mL in TBS (pH 7.5) using the Bradford assay (BioRad). Vaccine samples were aliquoted and stored at −80 °C until use. The concentration of HA in the final A/bald eagle/Florida/W22-134-OP/2022 (H5N1, 6:2 A/PR/8/34) vaccine was quantified in an indirect enzyme-linked immunosorbent assay (ELISA) using the 1H4 murine monoclonal antibody ^34^. Different dilutions of an H5 recombinant protein standard of known concentration were also included for absolute HA quantification.

### 4.5. SDS-PAGE and WB

An SDS-PAGE was performed to characterize the H5N1 inactivated split vaccine. Before running the SDS-PAGE, samples were deglycosylated with rapid PNGase F (New England Biolabs) according to manufacturer’s instructions. After deglycosylation, 20 μL of sample was mixed with 4X Laemmli buffer (BioRad) containing 50 mM NuPAGE sample reducing agent (dithiothreitol, Thermo Fisher Scientific). Samples were then incubated at 90 – 95 °C for 10 – 15 min and run on a 4 – 20% precast polyacrylamide Mini-PROTEAN TGX gel (BioRad) at 100 V for 10 min followed by 180 V for 35 min (30 μL/well). The gel was stained with InstantBlue Coomassie protein stain solution (Abcam) for 0.25 h before visualization. Images were taken in a Chemidoc MP Imaging System using the Image Lab software (BioRad).

For WB, SDS-PAGE gels were transferred to nitrocellulose membranes using an iBlot 2 transfer device (Invitrogen) at 25 V for 7 min. After, membranes were blocked in 3% (w/v) milk powder in PBS containing 0.1 % v/v Tween 20 (PBS-T) blocking solution at 4 °C in mild rocking conditions O/N. Membranes were then washed three times in PBS-T for 5 min, and an anti-H5 or an anti-N1 monoclonal antibody cocktails were added at ∼10 μg/mL and incubated at RT for 1 – 2 h. After washing three times in PBS-T for 5 min, horseradish peroxidase (HRP)-conjugated secondary anti IgG (H + L) polyclonal antibody (Invitrogen) was added at a 1:10000 dilution at RT for 1 h. Blots were again washed with PBS-T and developed by adding KPL TrueBlue peroxidase substrate (Seracare). Images were taken as previously described.

### 4.6. ELISA

Immulon 4 hBX 96-well plates (Thermo Fisher Scientific) were coated with 2 μg/mL of recombinant protein (50 μL per well) in PBS (pH 7.4) at 4 °C O/N. The next day, plates were washed three times with PBS-T and blocked in blocking solution (3 % v/v goat serum, 0.5 % w/v non-fat dry milk in PBS-T) for 1 h at RT. After blocking, mouse serum was added to the first well at a 1:30 dilution (150 μL/well) and serially diluted 1:3 in blocking solution and incubated for 2 h at 20 °C. Plates were washed three times with PBS-T before adding the secondary antibody (100 μL/well). For total IgG quantification, a 1:3000 dilution of sheep anti-mouse IgG (H&L) peroxidase conjugated (Rockland) in blocking solution was added. For IgG1 and IgG2a quantification, a 1:20,000 and 1:2,000 dilution in blocking solution of rabbit anti-mouse IgG1 or rabbit anti-mouse IgG2a (Invitrogen) was added, respectively. Afterwards, plates were incubated for 1 h at 20 °C and then washed four times with PBS-T with shaking. To develop plates, 100 μL of *O*-phenylenediamine dihydrochloride (OPD) substrate (SigmaFast OPD, Millipore Sigma) was added to each well. After a 10 min incubation, the reaction was stopped by adding 50 μL of 3M hydrochloric acid (HCl) to each well. The optical density at 490 nm (OD_490_) was measured on a Synergy H1 microplate reader (BioTek). A cut-off value of the average of the OD_490_ values of blank wells plus 3 times the standard deviation (SD) was established for each plate and used for calculating the area under the curve (AUC). AUC values were determined using GraphPad Prism 9 software.

### 4.7. ADCC reporter assay

ADCC activity in mouse sera was assessed using an FcγRIV cell-based ADCC reporter assay according to the manufacturer’s instructions (Promega). Briefly, white 96-well plates (Corning) were seeded with 2 × 10^4^ cells of MDCK cells per well and incubated O/N at 37 °C and 5% CO_2_. After 24 h, MDCK cells were washed with PBS and infected with A/bald eagle/FL/W22-134-OP/2022 (H5N1, 6:2 A/PR/8/34) or A/Vietnam/1203/2004 (H5N1, 6:2 A/PR/8/34) viruses at a multiplicity of infection of 5 at 37 °C and 5% CO_2_ for 1 h. After this, virus medium was aspirated, replaced with warm MEM, and incubated overnight at 37 °C and 5% CO_2_. The following day, the cell culture medium was removed and 25 μL of assay buffer [Roswell Park Memorial Institute (RPMI) 1640 medium supplemented with 4% v/v low IgG FBS, Gibco] was added to each well. Mouse sera previously heat-inactivated at 56 °C for 1 h were serially diluted 2-fold in RPMI 1640 medium and added to the infected MDCK cells (25 μL/well). The sera were incubated with MDCK cells at 37 °C for 30 min. Then, 7.5 × 10^4^ Jurkat cells expressing the mouse

FcγRIV with a luciferase reporter gene under transcriptional control of the nuclear factor-activated T cell promoter were added per well (25 μL/well) and incubated at 37 °C for 6 h. After incubation, 75 μL of Bio-Glo luciferase assay reagent was added per well and incubated at RT in the dark for 10 min. The luminescence signal was measured using a Synergy H1 microplate reader. The fold induction was calculated as follows: (RLU_induced_-RLU_background_)/(RLU_no antibody control_-RLU_background_), where RLU is relative luminescence units. The AUC values of the resulting fold-induction values were calculated using GraphPad Prism 10.

### 4.8. Neuraminidase inhibition assay

The NA activity of A/bald eagle/Florida/W22-134-OP/2022 (H5N1, 6:2 A/PR/8/34), A/Vietnam/1203/2004 (H5N1, 6:2 A/PR/8/34) and H7N1_A/Michigan/45/2015_ viruses was assessed on Immulon 4 hBX 96-well plates coated with 100 μL of fetuin (Millipore Sigma) at 25 μg/mL in PBS at 4 °C O/N. Fetuin-coated plates were washed three times with PBS-T and blocked with PBS + 5% v/v bovine serum albumin (BSA, MP Biomedicals). On a separate plate, the virus was serially diluted 1:2 in PBS + 1% w/v BSA, and 75 μL of pre-diluted virus samples were added to fetuin-coated plates already containing 75 μL of PBS + 1% w/v BSA. The fetuin-coated plates were incubated at 37 °C O/N. Afterwards, plates were washed four times with PBS-T with shaking, and 100 μL per well of peroxidase labeled peanut agglutinin from *Arachis hypogaea* (Millipore Sigma) at 5 μg/mL in PBS + 1% v/v BSA was added to the plates. Plates were incubated at 20 °C for 1.5 h before washing four times with PBS-T with shaking. To develop the plates, 100 μL of OPD substrate were added per well, incubated for 10 min at RT, and the reaction was stopped by adding 50 μL of 3M HCl per well. The OD_490_ was measured on a Synergy H1 microplate reader, and the half maximal effective concentration (EC_50_) was determined using the GraphPad Prism 9 software.

For the NAI assay, Immulon 4 hBX 96-well plates were coated with 100 μL of fetuin at 25 μg/mL in PBS at 4 °C O/N. Fetuin-coated plates were washed three times with PBS-T and blocked with PBS + 5% v/v BSA. In parallel, heat-inactivated sera at 56 °C for 1 h were serially diluted 1:2 in PBS + 1% v/v BSA with a starting dilution of 1:30 in non-fetuin coated 96-well plates (75 μL/well). Then, 75 μL of each virus corresponding to 2× EC_50_ was added per well to the pre-diluted sera plates and incubated at 20 °C for 1.5 h. After incubation, 100 μL of the virus/serum mixture were transferred per well to fetuin coated plates and incubated at 37 °C O/N. After incubation, the rest of the assay was performed as described above for the NA assay. No serum (virus only) and background controls (PBS + 1% v/v BSA only) were also included to measure the NAI. OD_490_ was measured on a Synergy H1 microplate reader, and the half-maximal inhibitory concentration (IC_50_) was calculated as: 1 – (OD_measured_-OD_background_)/(OD_no serum control_-OD_background_) in GraphPad Prism 10.

### 4.9. Microneutralization assay

Mouse sera were treated with receptor destroying enzyme (RDE) II (Denka Seiken) and incubated in a 37 °C water bath for 18-20 h. The same day, MDCK cells were seeded in 96-well cell-culture treated plates (Corning) at 1.8 × 10^4^ cells per well (100 μL/well) and incubated at 37 °C with 5% CO_2_ O/N. The following day, the RDE activity was stopped by the addition of a 2.5% w/v sodium citrate solution and incubation at 56 °C for 1 h. RDE-treated sera were initially diluted 1:10 and serially diluted 1:2 in infection medium consisting of minimum essential medium (MEM) with 10 mM of 4-(2-hydroxyethyl)-1-piperazineethanesulfonic acid (HEPES), 2 mM L-glutamine (Gibco), 3.2% w/v sodium bicarbonate (Corning), 1.2% w/v BSA, 100 U/mL penicillin and 100 μg/mL streptomycin (Gibco). Next, 120 μL of 100 × tissue culture infectious dose (TCID_50_) of virus prepared in infection medium and 120 μL of serially diluted sera were incubated on a shaker at RT for 1 h. MDCK cells were washed with 220 μL of PBS and incubated with 100 μL of the incubated serum-virus mixture at 37 °C with 5% CO_2_ for 1 h. Afterwards, the virus inoculum was carefully aspirated, MDCK cells were washed with PBS, and 100 μL of the serially-diluted sera containing 1 μg/mL of N-tosyl-L-phenylalanine chloromethyl ketone (TPCK)-treated trypsin (Millipore Sigma) were added to the cells and incubated at 37 °C with 5% CO_2_ for 48 h. As readout, the presence of virus was assessed by hemagglutination assay. In brief, 50 μL of cell supernatant was added to 96-well V-bottom plates (Nunc) and serially diluted 1:2. Then, 50 μL of 0.5% v/v turkey red blood cells (RBCs, Lampire Biological Laboratories) in PBS were added to each well, and plates were incubated on ice or at 4 °C for 45 min. The HA titer was calculated as the endpoint titer at which no RBC tear drop formation could be detected.

### 4.10. Hemagglutinin inhibition assay

A hemagglutination assay was initially performed to determine the hemagglutination titer units (HAU) of the viruses. Mouse sera were treated with RDE II and incubated in a 37 °C water bath for 18-20 h. The following day, the RDE activity was stopped by the addition of a 2.5% w/v sodium citrate solution and incubation at 56 °C for 1 h. RDE-treated sera were initially diluted 1:10 and serially diluted 1:2 in PBS in V-bottom 96-well plates. Twenty-five μl of serum dilutions were incubated with 25 μl of viruses diluted to 8 HAU at RT for 1 h. Following this, 50 μl of 0.5% v/v turkey RBCs diluted in PBS was added to the wells and incubated at 4 °C for 45 min before analysis as previously described.

### 4.11. Animal studies

All animal experiments were performed under protocols approved by the Icahn School of Medicine at Mount Sinai Institutional Animal Care and Use Committee. For all animal experiments conducted, 6–8-week-old female BALB/c (Jackson Laboratories) were used unless otherwise mentioned. The CpG 1018 (CpG) adjuvant (Dynavax Technologies) was used at 10 μg/mL, and the aluminum (Alum) hydroxide gel adjuvant (Alhydrogel 2%, InvivoGen) at 50 μg/mL. Mice were vaccinated via the intramuscular route with 0.0015 – 1.5 μg HA of H5N1 split vaccine prepared in Tris saline solution (20 mM Tris, 100 mM NaCl, pH 7.5) in a volume of 50 μL with or without adjuvants. A negative control (saline) consisting of Alum and CpG in Tris saline solution was also included. The vaccination regimen consisted of two sequential vaccinations 4 weeks apart. Four to six weeks after vaccination, mice were anesthetized and intranasally infected with 50 μL of influenza virus containing 0.1×, 5×, or 25×LD_50_ depending on the experiment. Additionally, mice were bled via submandibular bleeding for serological analysis at this time point. Blood was incubated at RT for 1 h and centrifuged at 5000 ×*g* for 30 min. Serum was separated from the pellet and stored at 4 °C until analysis. After virus challenge, weight loss was monitored for 14 days and mice showing a weight loss of ≥ 25% as compared with their initial body weight were humanely euthanized.

For the duration of the experiments, mice were housed in individually ventilated cages on a 12 h dark/light cycle with controlled temperature/humidity. Food and water were provided *ad libitum*.

### 4.12. Lung titers

Virus titers in the lungs of mice challenged with 0.5×LD_50_ of the A/bald eagle/Florida/W22-134-OP/2022 (H5N1, 6:2 A/PR/8/34) virus at days 3 and 6 after challenge were analyzed by the plaque assay method. Briefly, harvested lungs were homogenized in 2 disruption cycles (10 seconds/cycle) using tubes that contained high impact zirconium beads (Andwin Scientific) and 1 mL of PBS. For the plaque assay, lung homogenates were serially diluted 1:10 in PBS. Samples were incubated for 1 h with MDCK cells seeded at 3 × 10^5^ cells per well (1 mL/well) the day before in 12-well plates. After the 1 h incubation, an agarose overlay containing a final concentration of 0.64% w/v agarose (Oxoid) in MEM supplemented with 2 mM L-glutamine, 0.1% w/v of sodium bicarbonate, 10 mM HEPES, 100 U/mL penicillin, 100 μg/mL streptomycin, 0.2% w/v BSA, 1 μg/mL TPCK-treated trypsin, and 0.1% w/v diethylaminoethyl-dextran was added to the cells. The cells were incubated at 37 °C for 48 h, and visible plaques were counted after fixation with 3.7% v/v formaldehyde in PBS and visualization by immunostaining. All virus titers are presented as plaque forming units (PFU)/mL.

### 4.13. T-cell analysis

Mice were euthanized, the chest opened, and the right ventricle of the heart perfused with 10 mL of cold PBS. Lungs were processed using gentleMACS™ Tissue Dissociator (Miltenyi Biotech) and incubated in a collagenase-I (150 U/mL, Gibco)/DNAse-I (50 U/mL, Millipore Sigma) solution on a 37 °C water bath for 30 min. Following digestion, tissue was filtered through a 70 μm cell strainer (BD Biosciences). Spleens were mechanically disrupted with pestle homogenizers for 1.5 mL tubes (Fisher Scientific) and filtered as described above. Erythrocytes were lysed with RBC lysis buffer (BioLegend) in accordance with the manufacturer’s instructions. Cells were washed with 2% v/v FBS in PBS and reconstituted in 5 mL of complete media (RPMI 1640 supplemented with 10% v/v FBS, 100 U/mL penicillin, and 100 μg/mL streptomycin). Cells were counted in C-Chip™ Disposable Hemacytometers (SKC) using a 0.4% v/v Trypan blue solution to discriminate between live and dead cells. 2×10^6^ live cells per well were seeded in 96-well cell culture treated plates. For antigen-specific T-cellular immune response analysis, cells were stimulated with NP peptide pool PepTivator® Influenza A (H1N1) NP (Miltenyi Biotec) at 5 μg/mL, or with recombinant H5 and N1 glycoproteins (5 μg/mL) from A/bald eagle/Florida/W22-134-OP/2022 (H5N1). Stimulation with the NP peptide pool was performed at 37 °C with 5% CO_2_ for 6 h in the presence of 5 μg/mL Brefeldin A (BioLegend), 2 μM Monensin (BioLegend), and 25 μg/mL of co-stimulatory anti-CD28 antibodies (BioLegend). Stimulation with the H5 and N1 glycoproteins was performed at 37 °C and 5% CO_2_ for 18 h, with 5 μg/mL Brefeldin A, 2 μM Monensin, and 25 μg/mL anti-CD28 antibodies added 8 h after initial stimulation with the glycoproteins.

Following stimulation, cells were stained with fluorescently labeled antibodies (BioLegend) targeting different surface markers for the discrimination of naïve, effector memory, and central memory T-lymphocyte subpopulations (0.5 μL/sample CD3-BV711, 0.125 μL/sample CD4-PerCP/Cy5.5, 0.25 μL/sample CD8-BV785, 0.125 μL/sample CD62L-APC/Cy7, 0.25 μL/sample CD44-PE/Cy7). Intracellular staining of cytokines was performed with Cytofix/Cytoperm Fixation/Permeabilization Solution Kit (BD Biosciences) according to the manufacturer’s instructions. TNFα-AF488 (0.25 μL/sample), IFNγ-BV421 (1 μL/sample), and IL-2-PE (0.5 μL/sample) antibodies (BioLegend) were used to identify the corresponding cytokines. Data were collected on an Attune flow cytometer (Thermo Fisher Scientific) and analyzed with the FlowJo_v10.8.1 software. To calculate statistical differences between groups, the percentage of cytokine-producing cells in non-stimulated samples was subtracted from the corresponding values for peptide-stimulated samples.

### 4.14. B-cell enzyme-linked immunosorbent spot (ELISpot)

After euthanizing the mice, femurs and tibias were aseptically harvested, cleaned of muscle tissue, both ends of each bone cut, and placed in cold RPMI 1640 medium containing 2% v/v FBS. Two Eppendorf tubes, the one inside the other with a small hole on the bottom, was used to collect bone marrow cells after centrifugation at 1300 rpm for 7 min. The resulting cell suspension was gently pipetted up and down to disperse cell clumps and passed through a 70 μm cell strainer to obtain a single-cell suspension. RBCs were lysed using RBC lysis buffer, washed twice with PBS containing 2% v/v FBS, and resuspended in complete RPMI 1640 medium supplemented with 10% v/v FBS, 100 U/mL penicillin, and 100 μg/mL streptomycin. Cell viability and concentration were assessed by Trypan blue exclusion using a hemocytometer ^35^.

MultiScreen-IP plates (Millipore) were pre-wetted with 20 μL of 35% v/v ethanol, washed with water for injection (Gibco), and coated with 25 μg/mL of recombinant antigen in PBS (100 μL/well) at 4 °C O/N. The following day, plates were washed with PBS and blocked with RPMI 1640 containing 10% v/v FBS for at 37 °C for 1 h. After blocking, single-cell suspension obtained as described in the previous section were seeded at 2 × 10^5^ cells per well and incubated at 37 °C with 5% CO_2_ for 18 – 20 h to allow antigen-specific antibody secretion. After incubation, plates were washed thoroughly with PBS-T to remove cells and unbound antibodies. Biotinylated anti-mouse IgG detection antibody diluted in PBS with 0.5% v/v FBS was added to each well and incubated for at RT for 2 h. Plates were washed again with PBS, followed by incubation with streptavidin-alkaline phosphatase diluted in PBS with 0.5% v/v FBS at RT for 1 h. After a final wash, spots were developed using 5-Bromo-4-chloro-3-indolyl phosphate/Nitro blue tetrazolium (BCIP/NBT) substrate solution until distinct spots appeared, typically within 10 – 20 min. The reaction was stopped by rinsing plates extensively with tap water, and plates were air-dried overnight in the dark. Spots representing individual antibody-secreting B cells were imaged using an automated ELISpot reader (ImmunoSpot) and quantified using ImageJ. Results were expressed as the number of spot-forming cells per 10^6^ bone marrow cells.

### 4.15. Cytokine analysis

A mouse LEGENDplex kit (Biolegend) was used to measure the concentration of proinflammatory and Th1/Th2/Th17-associated cytokines in lung homogenates following the manufacturer’s instructions. Data were collected using an Attune flow cytometer and processed with LEGENDplex™ online data analysis software (legendplex.qognit.com).

### 4.16. nsEMPEM

Serum samples were heat-inactivated at 55°C for 1 h. For IgG isolation, 0.5 mL of mouse serum was mixed with 0.5 mL of pre-washed CaptureSelect IgG-Fc Affinity Matrix resin (Thermo Scientific) and 4 mL of PBS and incubated with gentle rotation at 4 °C for 48 h. The resin was washed three times with 5 mL of PBS, the IgG was eluted using 2.5 mL of 0.1 M Glycine (pH 2.0) into tubes containing 2 mL of 1 M Tris (pH 8) for immediate neutralization. For IgG digestion, papain was activated in 100 mM Tris-EDTA, 10 mM L-cysteine, and papain 1 mg/mL at 37 °C for 15 min. The activated papain was incubated with 1 mg of IgG at 37 °C for 5 h. The reaction was quenched with 0.05 M iodoacetamide. To purify fab, the digested mixture was incubated with CaptureSelect IgG-Fc affinity matrix rotating at 4°C for 1 h. The unbound supernatant containing the fab was collected, buffer-exchanged into Tris-buffered saline (TBS), and concentrated using 10 kDa Amicon ultra centrifugal filters (Merck Millipore).

Immune complexes were prepared in a molar proportion of ∼1:10 (antigen:fab), using 2 μg of antigen and 60 μg of polyclonal fab incubated at RT O/N. The unbound fab was washed using TBS in a 100 kDa Amicon ultra 0.5 mL centrifugal filter (Merck Millipore). The nsEMPEM grids were prepared using 3 μl of fab-antigen complexes in a concentration of 0.02 mg/mL and applied for 10 s to 400 mesh Cu^2+^ grids that were carbon coated, and glow discharged at 15 mA for 25 s. The fab-antigen complex was negatively stained with 2% v/v uranyl formate for 60 s. Data was collected using a Talos F200C electron microscope with a Ceta 16M camera at 200 kV and magnification of 73,000, pixel size of 2 Å/pixel. The defocus range was set between −2.5 and −2 mm and the electron dose was 27.1 e-/Å2. Automated data collection was carried out using EPU (Thermo Scientific).

After the automated data collection, Relion/4.0 was used for processing. Particles in the range of 190 to 280 Å were picked in the micrographs and 2D classification was performed using a box size of 208 pixels. Particles that contained trimer only or trimer-fab complexes were selected for 3D analysis. The 3D reference for 3D classifications and refinements was a low-resolution model of a non-liganded HA. Particles were then classified into 10 classes, and classes with similar features were combined and refined.

### 4.17. Statistical analysis

GraphPad Prism 10 was used for statistical analysis using Kruskal-Wallis and the post-hoc Dunn’s pairwise comparison test. The two-stage linear step-up false discovery rate procedure of Benjamini, Krieger, and Yekutieli was used to correct for multiple comparisons. Significance was considered with *p*-values equal or less than 0.05 (*), ≥0.01 (**), ≥0.001 (***), ≥0.0001 (****). The chosen sample size in each experiment is sufficient to generate statistically significant results.

## Data and materials availability

All data associated with this work can be found in the article, supplementary materials, and will be available via Immport. Negative stain electron microscopy maps representing each HA and NA epitope are deposited in the Electron Microscopy DataBank (EMDB) under accession IDs EMD-72062, EMD-72063, EMD-72064, EMD-72065, EMD-72066. All other data associated with this work can be found in the article or in supplementary materials. All data needed to evaluate the conclusions in the paper are present in the paper and/or the Supplementary Materials. Reagents and antigens described in the manuscript can be provided by the Krammer laboratory pending scientific review and a completed material transfer agreement (and any required shipping/handling permits for viruses). Requests for the reagents and antigens should be submitted to.

## Acknowledgements

We thank Maria Ibáñez-Trullén and Rajagowthamee R. Thangavel for technical support, and Ilaria Ceglia for project management (Department of Microbiology, Icahn School of Medicine at Mount Sinai, NY, USA). Some figure elements were obtained from NIH BioArt Source.

## Funding

This study was supported by the Collaborative Influenza Vaccine Innovation Centers (CIVIC) contract 75N93019C00051. Surveillance work that led to the detection of A/Canada goose/New York/NYCVH 22-9190/2022 used in this study was funded by Flu Lab and by a SEPA R25 (GM150146). Funding for generation of a sub-selection of reagents used was provided by Tito’s Vodka.

## Author contributions

E.P.-M. and F.K. conceived and designed the study. E.P.-M., T.E.G.A, M.J.S, K.V., H.A., and A.J.R. generated laboratory data. E.P.-M., T.E.G.A., K.V., H.A. and J.H. analyzed the data. J.Y., D.B., J.D.C., D.Y., R.J.W provided adjuvants, proteins, and viruses. E.P.-M., Y.K., G.N., J.H., A.B.W, and F.K. supervised the research. E.P.-M. wrote the original manuscript draft. All authors critically reviewed and approved the final version of the paper for submission.

## Competing interests

The Icahn School of Medicine at Mount Sinai has filed patent applications regarding influenza virus vaccines on which E.P.M. and F.K. are listed as inventors. F.K. has consulted for Merck, GSK, Gritstone, Sanofi, Curevac, Seqirus and Pfizer and is currently consulting for 3rd Rock Ventures and Avimex. The laboratory of F.K. is also collaborating with Dynavax on influenza vaccine development and with VIR on influenza virus therapeutics. A.B.W. has received royalty payments for the licensure of a prefusion coronavirus spike stabilization technology for which he is a co-inventor. A.B.W. and J.H. are currently consulting for Third Rock Ventures and Merida Biosciences. The laboratory of A.B.W. received unrelated sponsored research agreements from Third Rock Ventures during the conduct of the study. The authors declare that they have no other competing interests.

---

**Supplementary Figure S1.**
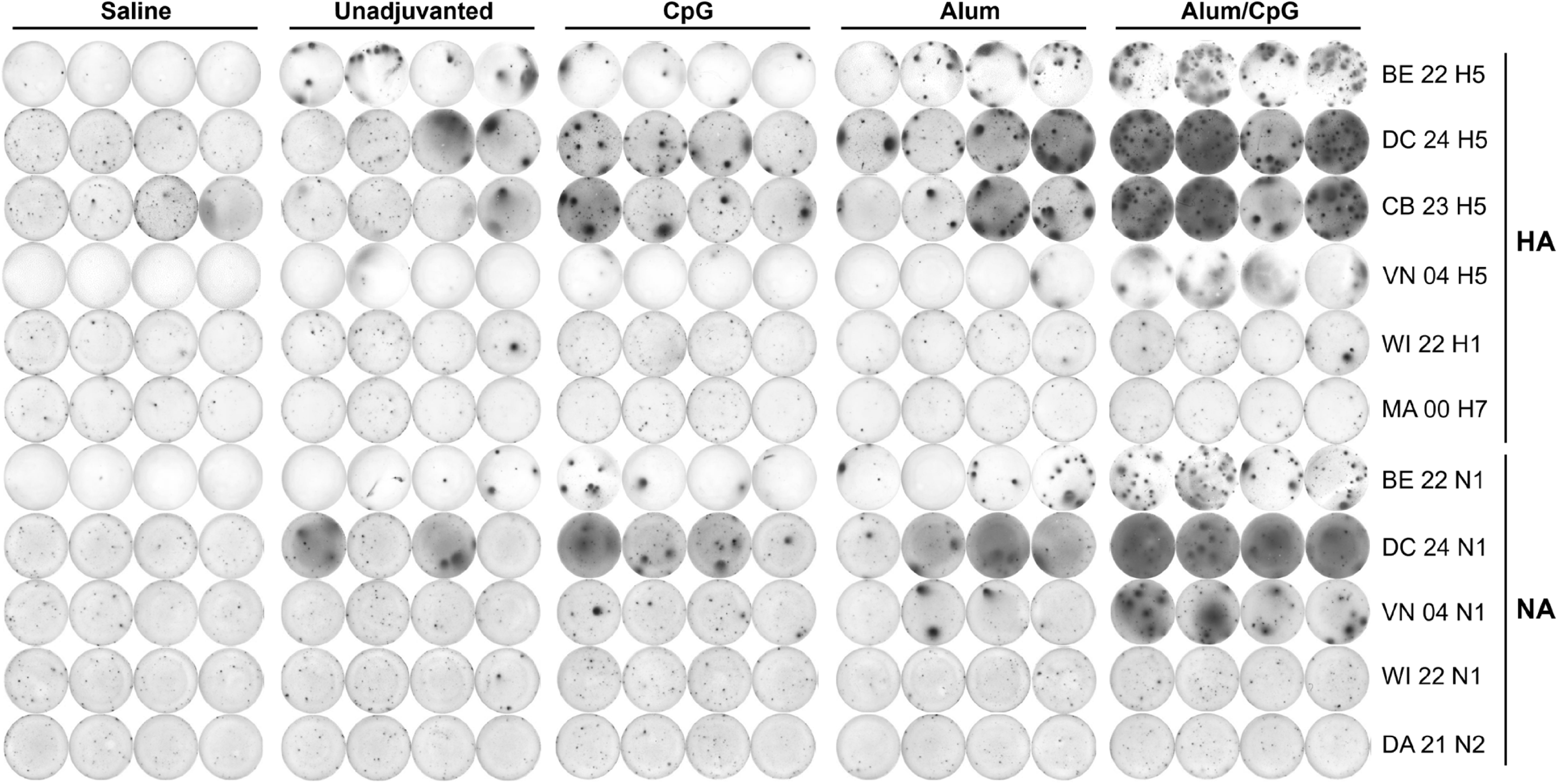
96-well ELISPOT plate used to quantify IgG antigen-specific secreting plasma cells in the bone marrow of different mice (*n* = 4) against different HA and NA glycoproteins. Bone marrow collection was conducted at 3 months post-vaccine prime. BE 22: A/bald eagle/FL/W22-134-OP/2022, DC 24: A/dairy cow/Texas/24-008749-001-original/2024, CB 23: A/Cambodia/NPH230032/2023, VN 04: A/Vietnam/1203/2004, WI 22: A/Wisconsin/67/2022, MA 00: A/mallard/Alberta/24/2001, DA 21: A/Darwin/6/2021.

**Supplementary Figure S2.**
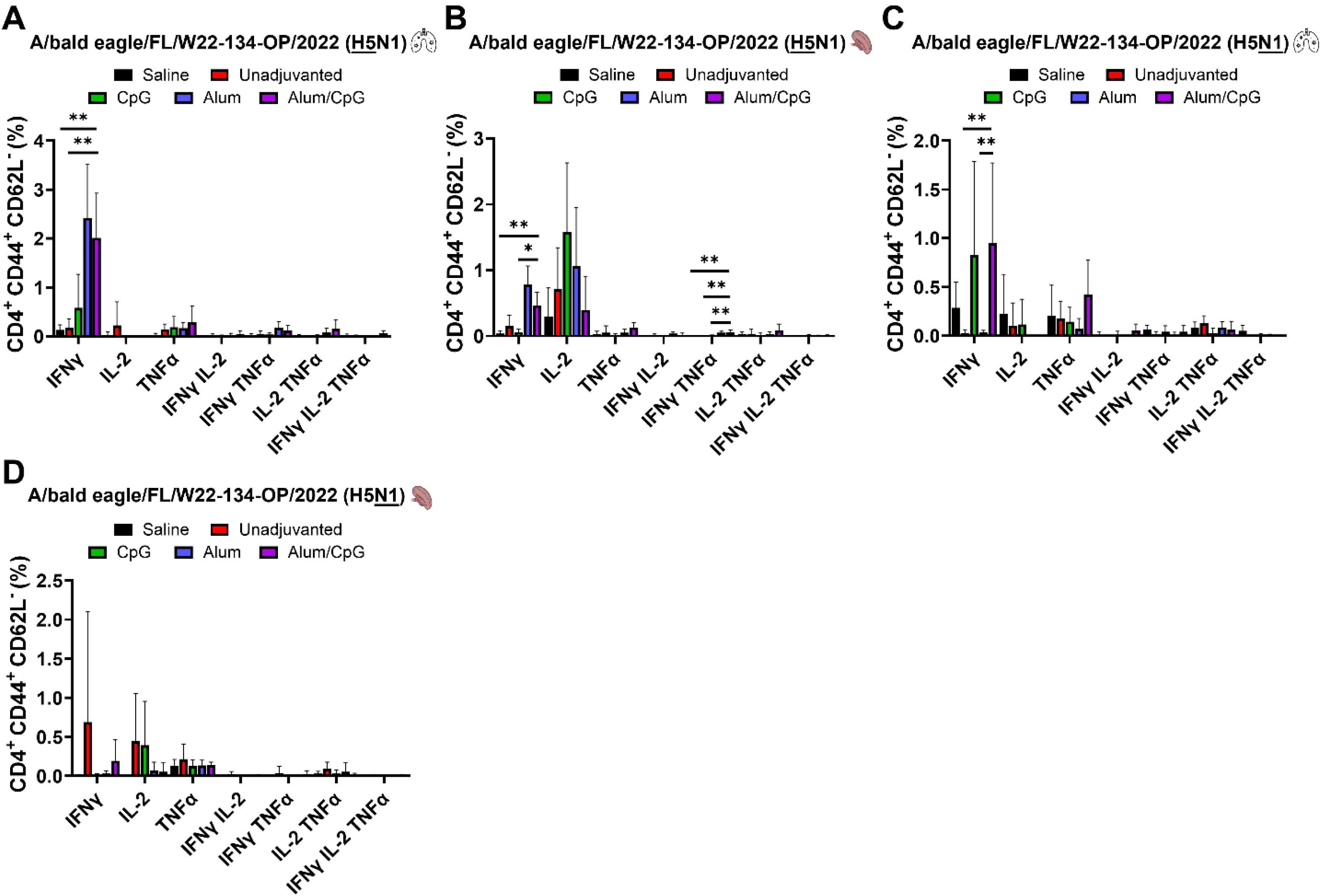
Percentage of IFNγ-, IL-2-, and TNFα-producing CD4^+^ effector memory (EM) T-lymphocytes in mouse spleens (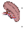) and lungs (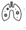) 5 days after challenge with 0.5×LD_50_ of A/bald eagle/Florida/W22-134-OP/2022 (H5N1, 6:2 A/PR/8/34) virus (*n* = 5) after background subtraction. T-cells were stimulated with H5 HA or N1 NA glycoproteins from A/bald eagle/Florida/W22-134-OP/2022. These different groups of mice were previously vaccinated in a prime/boost regimen with saline, 0.15 µg HA/mouse of split vaccine (unadjuvanted), and in combination with CpG, Alum, or Alum/CpG. Average and SD values are shown. Only statistically significant *p*-values (<0.05) are shown.

**Supplementary Figure S3.**
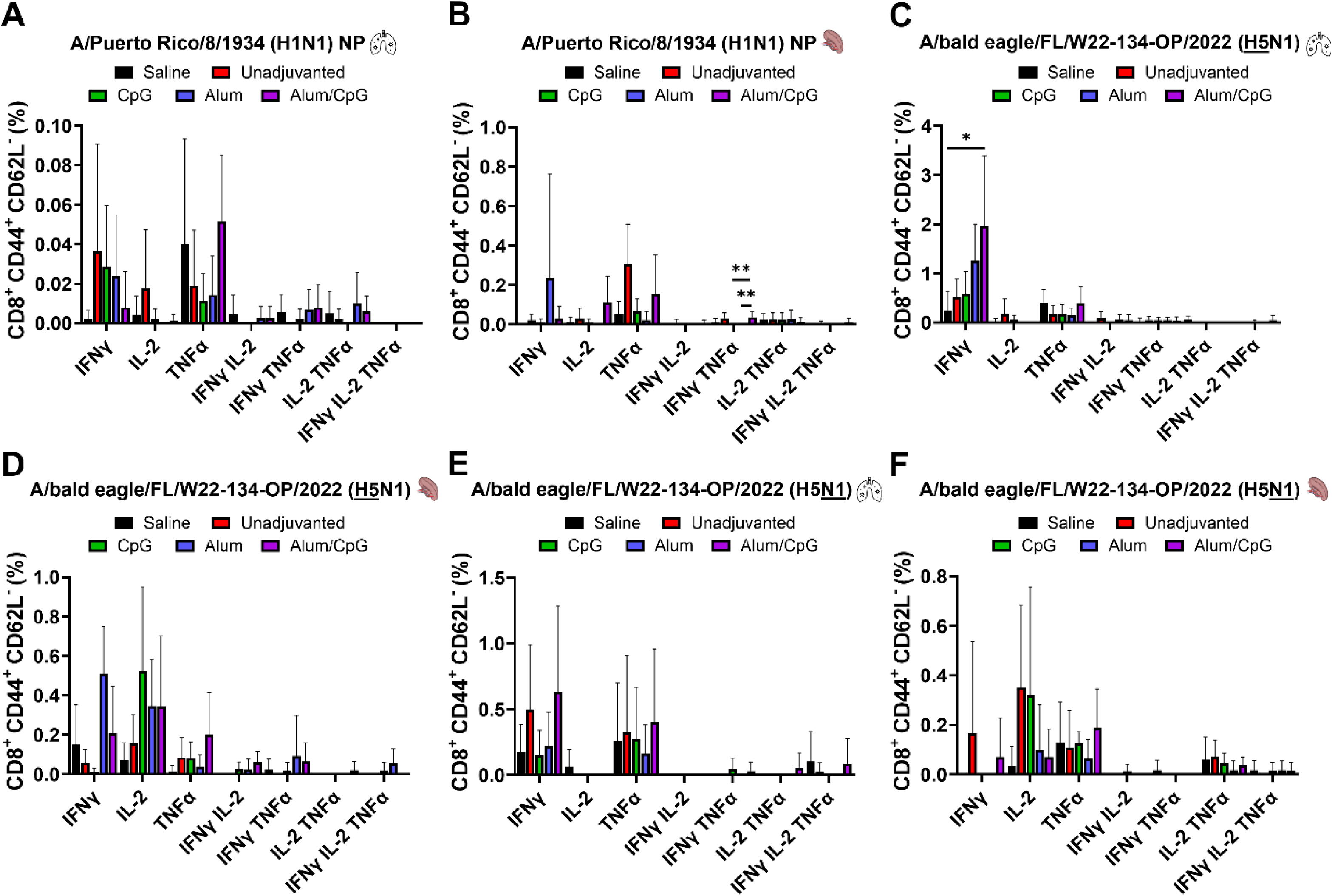
Percentage of IFNγ-, IL-2-, and TNFα-producing CD8^+^ effector memory (EM) T-lymphocytes in mouse spleens (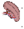) and lungs (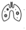) 5 days after challenge with 0.5×LD_50_ of A/bald eagle/Florida/W22-134-OP/2022 (H5N1, 6:2 A/PR/8/34) virus (*n* = 5) after background subtraction. T-cells were stimulated with H5 HA or N1 NA glycoproteins from A/bald eagle/Florida/W22-134-OP/2022 or with an ovelapping peptide library of the NP protein from A/Puerto Rico/8/1934 (H1N1) virus. These different groups of mice were previously vaccinated in a prime/boost regimen with saline, 0.15 µg HA/mouse of split vaccine (unadjuvanted), and in combination with CpG, Alum, or Alum/CpG. Average and SD values are shown. Only statistically significant *p*-values (<0.05) are shown.

**Supplementary Figure S4.**
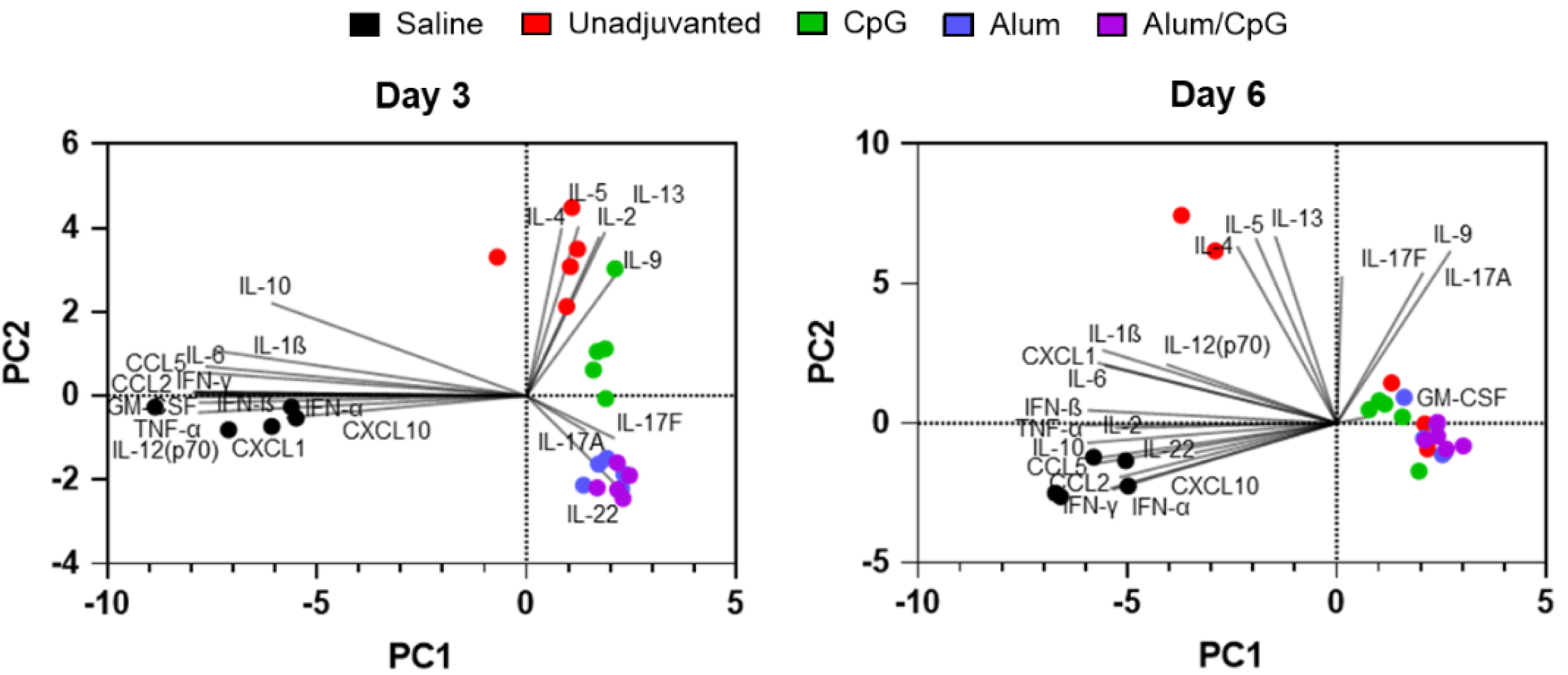
Principal component analysis (PCA) biplot representing the distribution of mice on the first two principal components (PC1 and PC2) based on cytokine responses in mouse lungs on day 3 (*n* = 5) and 6 (*n* = 5) post challenge with 0.5×LD50 of the A/bald eagle/Florida/W22-134-OP/2022 (H5N1, 6:2 A/PR/8/34) virus.

**Supplementary Figure S5.**
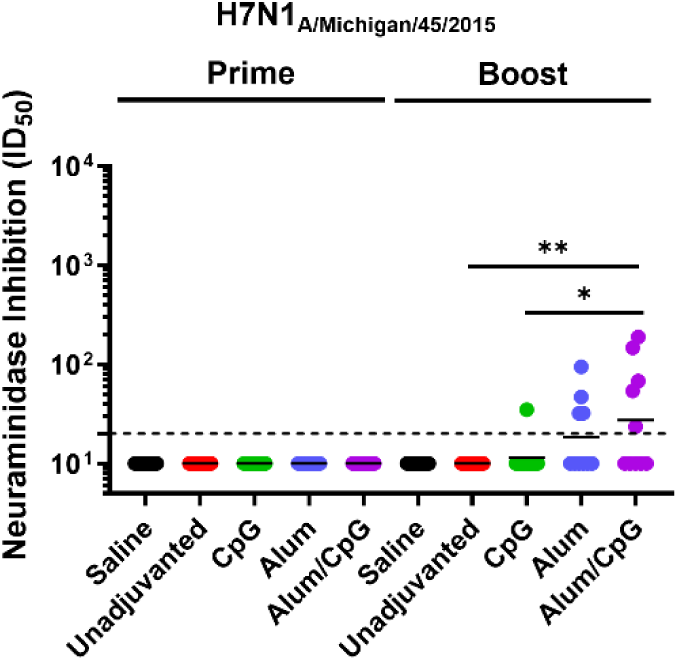
NAI assay of mouse sera (*n* = 10) against an H7N1 reassortant virus expressing the N1 of the A/Michigan/45/2015 (H1N1) virus. Individual and geometric mean ID_50_ serum dilution values are shown.

**Supplementary Figure S6.**
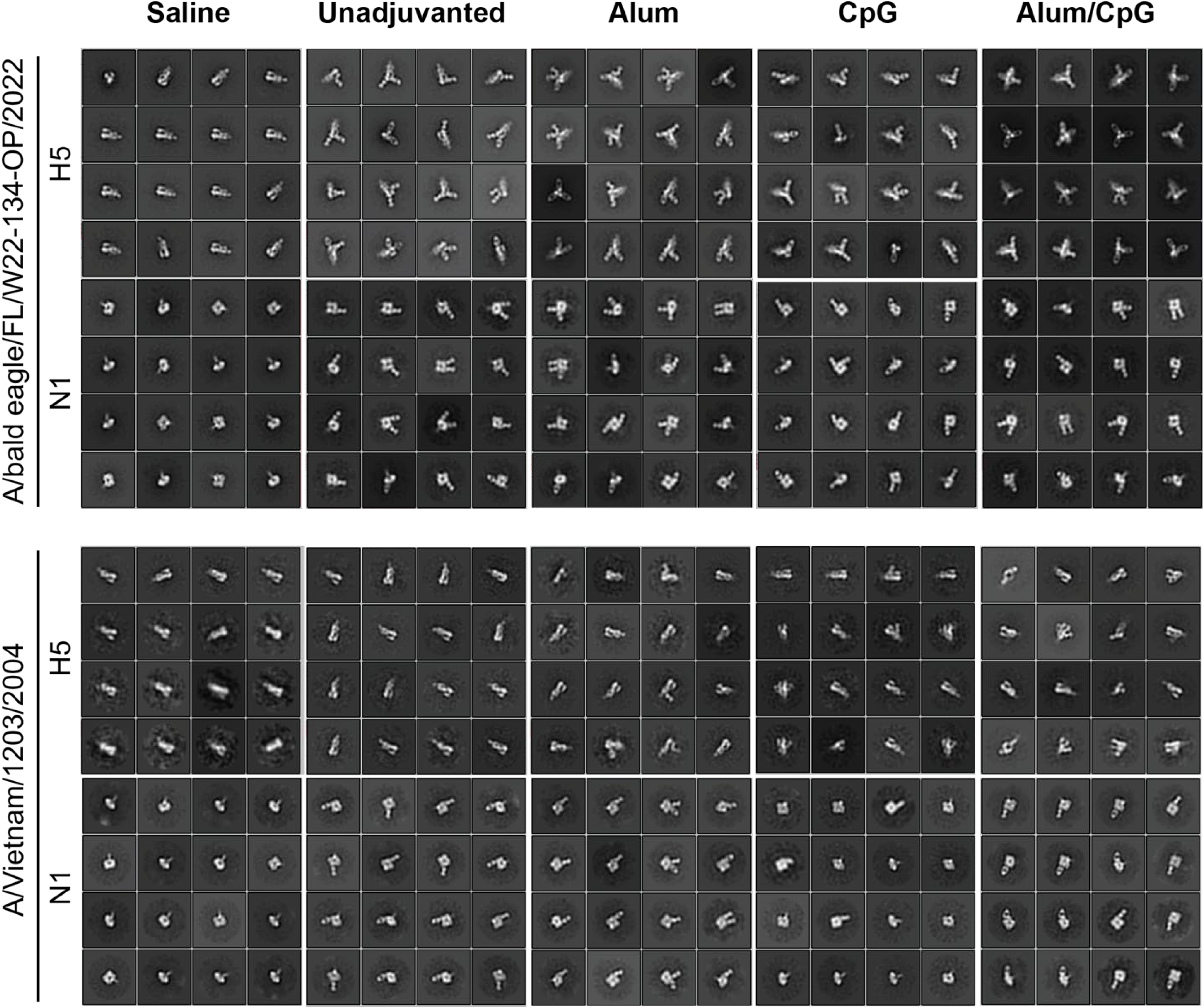
2D class averages from nsEMPEM of HAH5 BE22 and VN04 complexed with polyclonal fab isolated from mice by nsEMPEM after 3 – 4 weeks after second boost.

## References

1 Krammer, F., Hermann, E. & Rasmussen, A. L. Highly pathogenic avian influenza H5N1: history, current situation, and outlook. J Virol 99, e0220924 (2025). 10.1128/jvi.02209-24

2 Peacock, T. P. et al. The global H5N1 influenza panzootic in mammals. Nature 637, 304–313 (2025). 10.1038/s41586-024-08054-z

3 Taaffe, J. et al. An overview of influenza H5 vaccines. Lancet Respir Med 13, e20–e21 (2025). 10.1016/S2213-2600(25)00052-9

4 Khurana, S. et al. Licensed H5N1 vaccines generate cross-neutralizing antibodies against highly pathogenic H5N1 clade 2.3.4.4b influenza virus. Nat Med 30, 2771–2776 (2024). 10.1038/s41591-024-03189-y

5 Liedes, O. et al. Inactivated Zoonotic Influenza A(H5N8) Vaccine Induces Robust Antibody Responses Against Recent Highly Pathogenic Avian Influenza Clade 2.3.4.4b A(H5N1) Viruses. medRxiv, 2025.2002.2012.25322044 (2025). 10.1101/2025.02.12.25322044

6 Langley, J. M. et al. Safety and cross-reactive immunogenicity of candidate AS03-adjuvanted prepandemic H5N1 influenza vaccines: a randomized controlled phase 1/2 trial in adults. J Infect Dis 201, 1644–1653 (2010). 10.1086/652701

7 Ko, E. J. & Kang, S. M. Immunology and efficacy of MF59-adjuvanted vaccines. Hum Vaccin Immunother 14, 3041–3045 (2018). 10.1080/21645515.2018.1495301

8 Tzeng, T. T. et al. A TLR9 agonist synergistically enhances protective immunity induced by an Alum-adjuvanted H7N9 inactivated whole-virion vaccine. Emerg Microbes Infect 12, 2249130 (2023). 10.1080/22221751.2023.2249130

9 Puente-Massaguer, E. et al. Chimeric hemagglutinin split vaccines elicit broadly cross-reactive antibodies and protection against group 2 influenza viruses in mice. Sci Adv 9, eadi4753 (2023). 10.1126/sciadv.adi4753

10 Sicca, F., Neppelenbroek, S. & Huckriede, A. Effector mechanisms of influenza-specific antibodies: neutralization and beyond. Expert Rev Vaccines 17, 785–795 (2018). 10.1080/14760584.2018.1516553

11 Mosmann, T. R., McMichael, A. J., LeVert, A., McCauley, J. W. & Almond, J. W. Opportunities and challenges for T cell-based influenza vaccines. Nat Rev Immunol 24, 736–752 (2024). 10.1038/s41577-024-01030-8

12 Huang, X. et al. Increase in H5N1 vaccine antibodies confers cross-neutralization of highly pathogenic avian influenza H5N1. Nat Commun 16, 5517 (2025). 10.1038/s41467-025-60714-4

13 Puente-Massaguer, E. et al. Bioprocess development for universal influenza vaccines based on inactivated split chimeric and mosaic hemagglutinin viruses. Front Bioeng Biotechnol 11, 1097349 (2023). 10.3389/fbioe.2023.1097349

14 Zheng, X. et al. Alum/CpG adjuvant promotes immunogenicity of inactivated SARS-CoV-2 Omicron vaccine through enhanced humoral and cellular immunity. Virology 594, 110050 (2024). 10.1016/j.virol.2024.110050

15 Siegers, J. Y. et al. Emergence of a Novel Reassortant Clade 2.3.2.1c Avian Influenza A/H5N1 Virus Associated with Human Cases in Cambodia. medRxiv, 2024.2011.2004.24313747 (2025). 10.1101/2024.11.04.24313747

16 Krammer, F. Next-generation seasonal influenza virus vaccines need a neuraminidase component. Vaccine 54, 126994 (2025). 10.1016/j.vaccine.2025.126994

17 Hansen, L. et al. Human anti-N1 monoclonal antibodies elicited by pandemic H1N1 virus infection broadly inhibit HxN1 viruses in vitro and in vivo. Immunity 56, 1927–1938 e1928 (2023). 10.1016/j.immuni.2023.07.004

18 DiLillo, D. J., Palese, P., Wilson, P. C. & Ravetch, J. V. Broadly neutralizing anti-influenza antibodies require Fc receptor engagement for in vivo protection. J Clin Invest 126, 605–610 (2016). 10.1172/JCI84428

19 Sidney, J. et al. Targets of influenza human T-cell response are mostly conserved in H5N1. mBio 16, e0347924 (2025). 10.1128/mbio.03479-24

20 Hermann, E. & Krammer, F. Clade 2.3.4.4b H5N1 neuraminidase has a long stalk, which is in contrast to most highly pathogenic H5N1 viruses circulating between 2002 and 2020. mBio 16, e0398924 (2025). 10.1128/mbio.03989-24

21 Khurana, S. et al. AS03-adjuvanted H5N1 vaccine promotes antibody diversity and affinity maturation, NAI titers, cross-clade H5N1 neutralization, but not H1N1 cross-subtype neutralization. NPJ Vaccines 3, 40 (2018). 10.1038/s41541-018-0076-2

22 Paul, S. S. et al. A cross-clade H5N1 influenza A virus neutralizing monoclonal antibody binds to a novel epitope within the vestigial esterase domain of hemagglutinin. Antiviral Res 144, 299–310 (2017). 10.1016/j.antiviral.2017.06.012

23 Zhu, X. et al. A unique and conserved neutralization epitope in H5N1 influenza viruses identified by an antibody against the A/Goose/Guangdong/1/96 hemagglutinin. J Virol 87, 12619–12635 (2013). 10.1128/JVI.01577-13

24 Ekiert, D. C. et al. Antibody recognition of a highly conserved influenza virus epitope. Science 324, 246–251 (2009). 10.1126/science.1171491

25 Dreyfus, C. et al. in Science Vol. 337 1343-1348 (American Association for the Advancement of Science, 2012).

26 Wan, H. et al. Structural characterization of a protective epitope spanning A(H1N1)pdm09 influenza virus neuraminidase monomers. Nat Commun 6, 6114 (2015). 10.1038/ncomms7114

27 Hatta, M. et al. An influenza mRNA vaccine protects ferrets from lethal infection with highly pathogenic avian influenza A(H5N1) virus. Sci Transl Med 16, eads1273 (2024). 10.1126/scitranslmed.ads1273

28 Furey, C. et al. Development of a nucleoside-modified mRNA vaccine against clade 2.3.4.4b H5 highly pathogenic avian influenza virus. Nat Commun 15, 4350 (2024). 10.1038/s41467-024-48555-z

29 Margine, I., Palese, P. & Krammer, F. Expression of functional recombinant hemagglutinin and neuraminidase proteins from the novel H7N9 influenza virus using the baculovirus expression system. J Vis Exp, e51112 (2013). 10.3791/51112

30 Leon, A. N. et al. Structural mapping of polyclonal IgG responses to HA after influenza virus vaccination or infection. mBio 16, e0203024 (2025). 10.1128/mbio.02030-24

31 Kandeil, A. et al. Rapid evolution of A(H5N1) influenza viruses after intercontinental spread to North America. Nat Commun 14, 3082 (2023). 10.1038/s41467-023-38415-7

32 Guan, L. et al. Cow’s Milk Containing Avian Influenza A(H5N1) Virus - Heat Inactivation and Infectivity in Mice. N Engl J Med 391, 87–90 (2024). 10.1056/NEJMc2405495

33 Hai, R. et al. Influenza B virus NS1-truncated mutants: live-attenuated vaccine approach. J Virol 82, 10580–10590 (2008). 10.1128/JVI.01213-08

34 Rajendran, M. et al. An immuno-assay to quantify influenza virus hemagglutinin with correctly folded stalk domains in vaccine preparations. PLoS One 13, e0194830 (2018). 10.1371/journal.pone.0194830

35 Zhu, Y. P. et al. Preparation of Whole Bone Marrow for Mass Cytometry Analysis of Neutrophil-lineage Cells. J Vis Exp (2019). 10.3791/59617

36 Alzua, G. P. et al. Human monoclonal antibodies that target clade 2.3.4.4b H5N1 hemagglutinin. bioRxiv, 2025.2002.2021.639446 (2025). 10.1101/2025.02.21.639446

